# Proximity Analysis of Native Proteomes Reveals Interactomes Predictive of Phenotypic Modifiers of Autism and Related Neurodevelopmental Conditions

**DOI:** 10.1101/2022.10.06.511211

**Authors:** Yudong Gao, Matthew Trn, Daichi Shonai, Jieqing Zhao, Erik J. Soderblom, S. Alexandra Garcia-Moreno, Charles A. Gersbach, William C. Wetsel, Geraldine Dawson, Dmitry Velmeshev, Yong-hui Jiang, Laura G. Sloofman, Joseph D. Buxbaum, Scott H. Soderling

## Abstract

One of the main drivers of autism spectrum disorder is risk alleles within hundreds of genes, which may interact within shared but unknown protein complexes. Here we develop a scalable genome-editing-mediated approach to target 14 high-confidence autism risk genes within the mouse brain for proximity-based endogenous proteomics, achieving high specificity spatial interactomes compared to prior methods. The resulting native proximity interactomes are enriched for human genes dysregulated in the brain of autistic individuals and reveal unexpected and highly significant interactions with other lower-confidence autism risk gene products, positing new avenues to prioritize genetic risk. Importantly, the datasets are enriched for shared cellular functions and genetic interactions that may underlie the condition. We test this notion by spatial proteomics and CRISPR-based regulation of expression in two autism models, demonstrating functional interactions that modulate mechanisms of their dysregulation. Together, these results reveal native proteome networks *in vivo* relevant to autism, providing new inroads for understanding and manipulating the cellular drivers underpinning its etiology.

## Introduction

Autism spectrum disorder (hereinafter “autism”) is a neurodevelopmental condition associated with social communication difficulties and restricted and repetitive behaviors. Autism presents with significant clinical heterogeneity and complex genetic etiology ^1^. Decades of research have identified and curated an evolving list of gene mutations associated with autism risk, many of which converge on pathways mediating synaptic/axonal functions and gene regulation ^2–7^, and exhibit cell-type specific expression patterns across the brain ^8^. Until recently, evidence of shared biology across these risk genes has been inferred primarily from RNA-level gene expression results ^9–11^, or interpreted from protein interaction analyses ^12–15^, although these are often derived from non-neuronal cells. Thus, there is a prevailing notion in the literature that molecular convergence in autism may be optimally reflected at the protein level in the brain.

One hallmark of neurons is their distinctive sub-cellular compartmentation - such as the synapse and axonal initial segment (AIS) - that are pivotal for neurotransmission ^16,17^. In addition, gene expression regulators, many of which reside within the nucleus, also orchestrate key milestones of neurodevelopment ^6^. It is not surprising that autism risk converges on genes encoding proteins associated with these sub-cellular compartments ^18–22^. Identifying the autism-associated proteome architecture of these compartments could define how seemingly diverse genetic mutations are functionally connected, thereby providing a new roadmap to reveal a converging autism etiology. Current efforts to dissociate protein complexes and protein-protein interactions (PPIs) rely upon techniques such as cellular fractionation, immunoaffinity purification, cross-linking, and proximity-based proteomic methods, including biotin-identification (BioID) ^23–30^. However, due to inherent specificity constraints of antibody-dependent immunoprecipitation or recombinant exogenous promoter-based strategies that express proteins at non-physiologic levels in non-native cell types, access to the native proximity interactomes of endogenously expressed autism risk proteins within brain tissue remains a significant challenge. Given these limitations, high-fidelity proteomes organized around autism risk proteins in the brain are clearly needed for better mechanistic insights.

Previously, we published a two-vector CRISPR/Cas9 approach, Homology independent Universal Genome Engineering (HiUGE), which enables rapid endogenous neuronal gene knock-in for protein modification *in vivo* ^31^. Here, we have leveraged HiUGE and simplified it with a one-vector design to achieve a robust knock-in in brain tissue with an engineered biotin ligase, TurboID ^32^, for scalable proximity proteomics of endogenous protein complexes. The approach is flexible also from a protein engineering perspective, as the proteins can be modified by placing TurboID within terminal coding exons or internally within introns using splicing sequences ^33^. Importantly, because this strategy uses AAV transduction of readily available Cas9-transgenic mice, it obviates the significant time and cost burden of the traditional method of generating *in vivo* fusions of endogenous proteins by producing transgenic mice via germline transmission. Coupling this approach with *in vivo* BioID (iBioID) ^23^, we have targeted 14 genetic drivers of autism that are mapped to the synapse, AIS, or the nucleus. Out of these 14 targets, 13 are SFARI gene score 1, one is gene score 2, also noted as syndromic, and 7 are from the autism risk gene lists of Satterstrom et al., 2020 ^3^, or Fu et al., 2022 ^7^. With this approach, we have modified the endogenous proteins with TurboID and then unraveled their native proximal proteomes directly from brain tissue for the first time to our knowledge. We have identified 1252 proteins within these 14 proximity proteomic interactomes (hereinafter “interactomes”) and 3264 proximity PPIs associated with these autism targets. Amongst them, 16% are proteins encoded by mouse orthologs of SFARI genes ^2^, 8% overlap with differentially expressed genes (DEGs) found in brain tissue of autistic individuals ^8^, and 65% of the PPIs are not reported in STRING queries ^34,35^. Importantly, direct comparisons of HiUGE-iBioID to the prior “gold standard” of immunoprecipitation revealed its interactomes are more specific to the baits’ biological functions, containing far fewer proteins with potential off-target functions. Similar conclusions are drawn from comparison with results obtained from non-native recombinant BioID expression *in vitro*, supporting the advantages of native interactome discovery using HiUGE-iBioID *in vivo*. Notably, the interactomes contain products of many newly discovered autism risk genes identified in recent studies ^7,36–38^ and reveal shared biological processes among autism proteins that may be predictive of interactions influencing autism phenotypes.

We have tested this notion by identifying intersections between HiUGE-iBioID interactomes and co-perturbed proteins in two autism mouse models (Syngap1 synaptopathy and Scn2a channelopathy). In the Syngap1 model, we find that its binding with Anks1b is disrupted by an autism-associated Syngap1 mutation, and their interaction is essential for shaping neural activity during critical synaptogenesis periods. In the Scn2a model, we show that a patient-derived missense mutation results in repetitive behaviors and abnormal social communication in mice. The mutation also downregulates a key Scn2a modulatory protein cluster discovered in its interactome and results in aberrant attenuation of neural activity. Strikingly, re-expression of this cluster rescues this autism-associated electrophysiological impairment.

Together, our results establish a new scalable platform to engineer endogenous proteins and map native proximity interactomes at the protein-level that are associated with genetic risks for neurodevelopmental conditions such as autism. Our findings also reveal an intersectional approach to prioritize candidates based on proteomic co-perturbation. These data support a protein-centric model to reveal novel mechanisms of autism and related neurodevelopmental disorders and potential mitigation approaches.

## Results

### Endogenous labeling of autism risk proteins using HiUGE

Previously we have used overexpression of bait proteins fused to various biotin ligases to discover interactomes from brain tissue ^23,39^. Although this approach has been highly successful, it is known that the expression of proteins using artificial promoters may result in non-native interactions due to non-physiological expression levels and/or expression in inappropriate cell types. To overcome these issues, cell lines harboring CRISPR-edited knock-ins (KIs) of TurboID have been utilized ^40^. Nonetheless, such approaches are limited by the inability to recapitulate the diversity of neuronal cell types that exist *in vivo*, and cultured cells cannot replicate many conditions governed by native neuropil interactions. To address this technological gap, we developed a new approach for iBioID experiments with in-frame TurboID fusions introduced into endogenous gene regions by HiUGE genome editing (Fig. 1A). Using a highly expressed gene Tubb3 as a pilot example, we confirmed that the HiUGE-iBioID yields efficient *in vivo* KI and biotinylation in neurons across the brain (Fig. S1A).

**Fig. 1.**
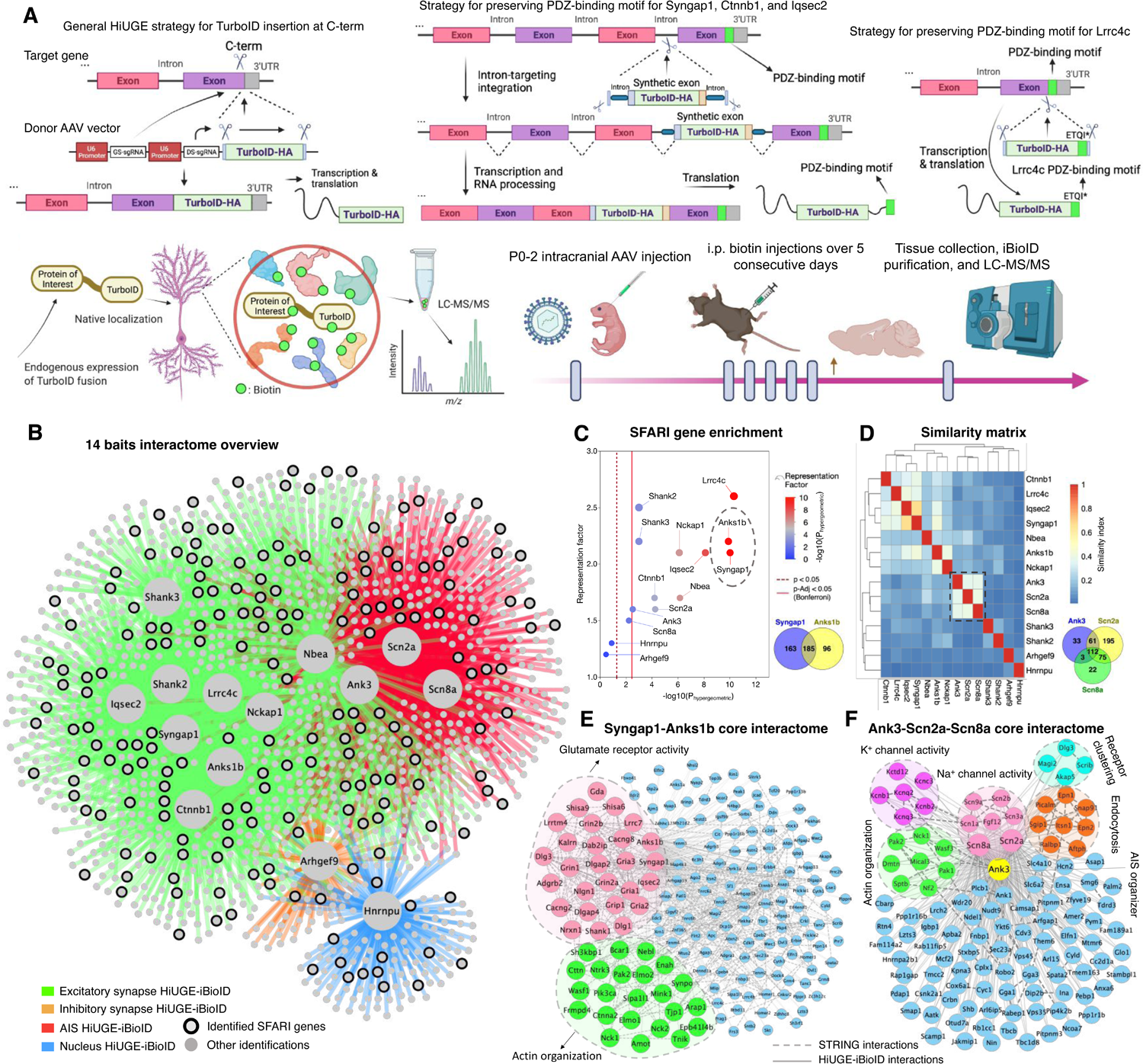
HiUGE-iBioID reveals endogenous interactomes of 14 autism risk proteins. (**A**) Schematic illustration of HiUGE-iBioID and its workflow. GS-sgRNA: gene-specific-gRNA. DS-sgRNA: donor-specific-gRNA. Strategies to preserve C-term PDZ-binding motifs of Syngap1, Ctnnb1, Iqsec2, and Lrrc4c are detailed. (**B**) Overview of 14 interactomes that segregates according to expected bait functions. (**C**) Enrichment analysis of overlapping SFARI genes using hypergeometric probability. (**D**) Interactome clustering based on a similarity matrix. (**E**) Core interactome between Syngap1 and Anks1b that show highly significant overlaps with SFARI genes. (**F**) Core interactome amongst Ank3, Scn2a, and Scn8a that are clustered based on similarity. Modules of proteins were isolated by MCL clustering or GO analysis.

Because autism is driven by mutations in a large number of risk genes whose products may interact with each other in unknown functional ways, we next targeted 14 high-confidence autism risk genes from the SFARI gene list (*Anks1b*, *Syngap1*, *Shank2*, *Shank3*, *Nckap1*, *Nbea*, *Ctnnb1*, *Lrrc4c*, *Iqsec2*, *Arhgef9*, *Ank3*, *Scn2a*, *Scn8a*, and *Hnrnpu*) that are expressed in neuronal compartments of the synapse, AIS, and nucleus ^2,41,42^. Note, due to packaging limits, many of these protein targets are too large to overexpress using conventional AAV methods, which further supports the need to label the endogenous copies of these genes. Importantly, for targets with C-terminal (C-term) PDZ-binding motifs (Syngap1, Ctnnb1, Iqsec2, and Lrrc4c), intron-targeting ^33^ or custom donor strategies were used to preserve these critical protein-interaction sites (Fig. 1A). First, to confirm the proper localization of HiUGE-labeled targets, a highly immunogenic spaghetti monster (smFP) tag ^43^, similar in size to TurboID, was used to visualize fusion proteins. We found that HiUGE-labeled proteins were either properly localized to the synaptic sites, colocalizing with the Homer1 immunosignal (Fig. S1B-J, Q), or were restricted to the distinct features of the AIS and nuclear compartments (Fig. S1L-O). We further validated that the HiUGE-labeling was colocalized with the immunofluorescence of specific antibodies, demonstrating the localization of these proteins was not affected by the tag fusion (Fig. S2). Having thus confirmed proper genome editing and correct fusion protein localization, we next fused each protein *in vivo* with TurboID-HA by injecting HiUGE AAV directly into Cas9 transgenic neonatal pup brains (P0-2), and then biotinylated surrounding proteins by supplying biotin via intraperitoneal (i.p.) injections over 5 consecutive days starting at ∼P21. Western blot analyses of the purified streptavidin-precipitations from forebrain lysates collected at ∼P26 detected epitope-tagged baits at the expected molecular masses, confirming correct TurboID fusion protein expression (Fig. S3A-H).

### HiUGE-iBioID reveals endogenous interactomes associated with autism risk proteins of diverse subcellular compartments

LC-MS/MS analysis of the 14 HiUGE-iBioID samples detected a total of 1252 proteins that were enriched in the bait proteomes, with expected interactions faithfully captured (Fig. 1B, S11-24, Table S1,2). Importantly, 65% of the interactions detected were new, being absent from STRING queries (using a generous stringency interaction score of 0.15 ^34,35^) - likely a reflection of prior studies being conducted in non-neuronal cell types or methods other than proximity proteomics. Gene ontology (GO) analyses revealed highly cohesive cellular functions corresponding to the known biology of the bait proteins, such as pathways associated with synaptic transmission (Anks1b, Syngap1, Shank2, Shank3, Nckap1, Nbea, Ctnnb1, Lrrc4c, Iqsec2, Arhgef9), voltage-gated channel activity (Ank3, Scn2a, Scn8a), and RNA processes (Hnrnpu); thereby demonstrating high fidelity identification of local interactomes (Fig. S11-24, S27). These networks were also consistent with the latest knowledge of the structures and functions of specific neuronal compartments. For example, the detection of Mical3 and Septin complexes with the AIS baits (Fig. S21-23, Table S2) echoed a previous study that suggested their roles in regulating cytoskeletal stability and polarized trafficking at the AIS ^24^. Reciprocal analyses also supported the reproducibility of the interactomes, showing that the baits frequently cross-identified each other (Fig. S4A, Table S2). Additional HiUGE-iBioID of proteomic “hubs” (e.g., Homer1 and Wasf1, which were commonly associated with many baits) demonstrated they could reversely detect the majority of baits in validation experiments (Fig. S4B-H). In addition, comparisons showed that HiUGE-iBioID interactomes significantly overlap with, but also differ from proteomic detections derived from independent antibody immunoprecipitations we performed for Anks1b and Scn2a (Fig. S4I, J). Interactions specific to HiUGE-iBioID compared to these antibody pulldowns were expected since BioID is based on covalent labeling of nearby proteins and thus does not require transient or weak interactions to survive the pulldown and washing steps of immunoaffinity purification. Of note, proteomic detections unique to HiUGE-iBioID conformed to the expected top molecular function GO terms of Anks1b and Scn2a, while detections unique to immunoprecipitations did not (Fig. S4K, L). These data demonstrate that HiUGE-iBioID excels in specificity for detecting biologically-relevant proteomic interactions.

We also sought to determine if the interactomes showed a significant overlap with differentially expressed genes (DEGs) from specific cell types in autistic individuals by cross-referencing a recent human tissue single-cell genomics study ^8^. HiUGE-iBioID networks showed the highest level of overlap with autism DEGs in cortical layer 2/3 (L2/3) excitatory neurons (Fig. S5A), consistent with the study suggesting L2/3 neurons are significantly affected in autism ^8^. Notably, most networks were also significantly enriched for other autism risk genes (SFARI, Satterstrom et al. ^3^, and Fu et al. ^7^; Fig. 1C, S5B), demonstrating the convergence of autism genetic susceptibilities at the protein interaction level in previously unknown ways. We also performed enrichment analyses between these networks and an autism genome-wide association meta-analysis (GWAS) study ^44^; however, the result was largely nonsignificant or marginal, likely due to limited power (Table S7). It has also been noted that larger autism cohorts ^38^, or new approarchs, are needed to determine the potential genome-wide significance of a large number of moderate-risk genes. Importantly, the HiUGE-iBioID networks of high-confidence autism risk proteins formed physical communities with other candidate proteins of moderate confidence (Table S2, overlap tab). These genes represent a resource of moderate autism candidates that likely should be prioritized in future studies of genetic contributions due to their possible functional interactions with high-confidence autism risk genes. Furthermore, we noted the interactomes contained numerous potentially druggable targets, including ∼ 80 kinases and phosphatases, 18 of which are encoded by SFARI gene mouse orthologs: Camk2a, Camk2b, Cask, Cdc42bpb, Cdkl5, Csnk1e, Csnk2a1, Dyrk1a, Mapk3, Ntrk2, Ntrk3, Ocrl, Pak1, Pak2, Ppp3ca, Rps6ka2, Taok2, Wnk3.

Finally, similarity clustering of the bait interactomes revealed subgroups that largely segregated according to their expected cellular compartments (Fig. 1D). Networks of the overlapping interactomes between two baits with highly significant SFARI gene enrichment (Syngap1 and Anks1b) and three similarity-clustered AIS baits (Ank3, Scn2a, and Scn8a) revealed shared pathways that emphasized glutamate receptor and voltage-gated channel activities, respectively, and common modules that were involved in actin organization in both networks (Fig. 1E-F). These interactomes may contain proteins that function together with the autism-associated baits, modulation of which may reveal the complex genetic risks of autism or serve as potential inroads for normalizing relevant phenotypes.

### Syngap1 mutations lead to reshaping of its synaptic interactome and loss of Anks1b binding

We hypothesized that protein interactome discovery could inform functional interactions relevant to the phenotypes underlying autism. To test this proposition, we first focused on the intersection between Syngap1 and Anks1b, as their proteomes show a high degree of overlap and exhibit significant SFARI gene enrichment (Fig. 1C). Markov cluster algorithm (MCL) ^45^ analysis of their overlapping interactome identified a large PPI community (27 proteins, including Syngap1 and Anks1b), that was significantly enriched for pathways of “autistic disorder” and “glutamate receptor activity”, indicating this shared cluster may regulate excitatory synaptic transmission in autism. Although molecular interaction between Syngap1-Anks1b has been reported previously ^27,46–48^, an understanding of their functional interaction remains limited. Prior studies have indicated that Anks1b and Syngap1 both regulate synaptic activity and plasticity through NMDA-type glutamate receptors ^49,50^, they are dispersed from the PSD in response to synaptic activity in a CaMKII-dependent manner ^51–53^, and are associated with similar autism-like phenotypes in haploinsufficiency models ^54,55^. Hence, we sought to test whether Syngap1 and Anks1b functionally interact in driving neuronal phenotypes.

We first analyzed the synaptic proteomes in wildtype (WT) and Syngap1 heterozygous (Syngap1-Het) mice ^56–58^ (Fig. 2A) to determine if Anks1b was influenced by haploinsufficiency of Syngap1. Fractionation steps were performed to isolate the synaptosomes ^26,59,60^ from the cerebral cortex and striatum. In Syngap1-Het mice, Anks1b was depleted (∼60%) to a level similar as for Syngap1 in both cortical and striatal synaptosomes of Syngap1-Het mice (Fig. 2B, Table S4), supporting the notion of a very close functional interaction between Syngap1 and Anks1b not only in typical physiology, but also in autism-associated synaptopathy.

**Fig. 2.**
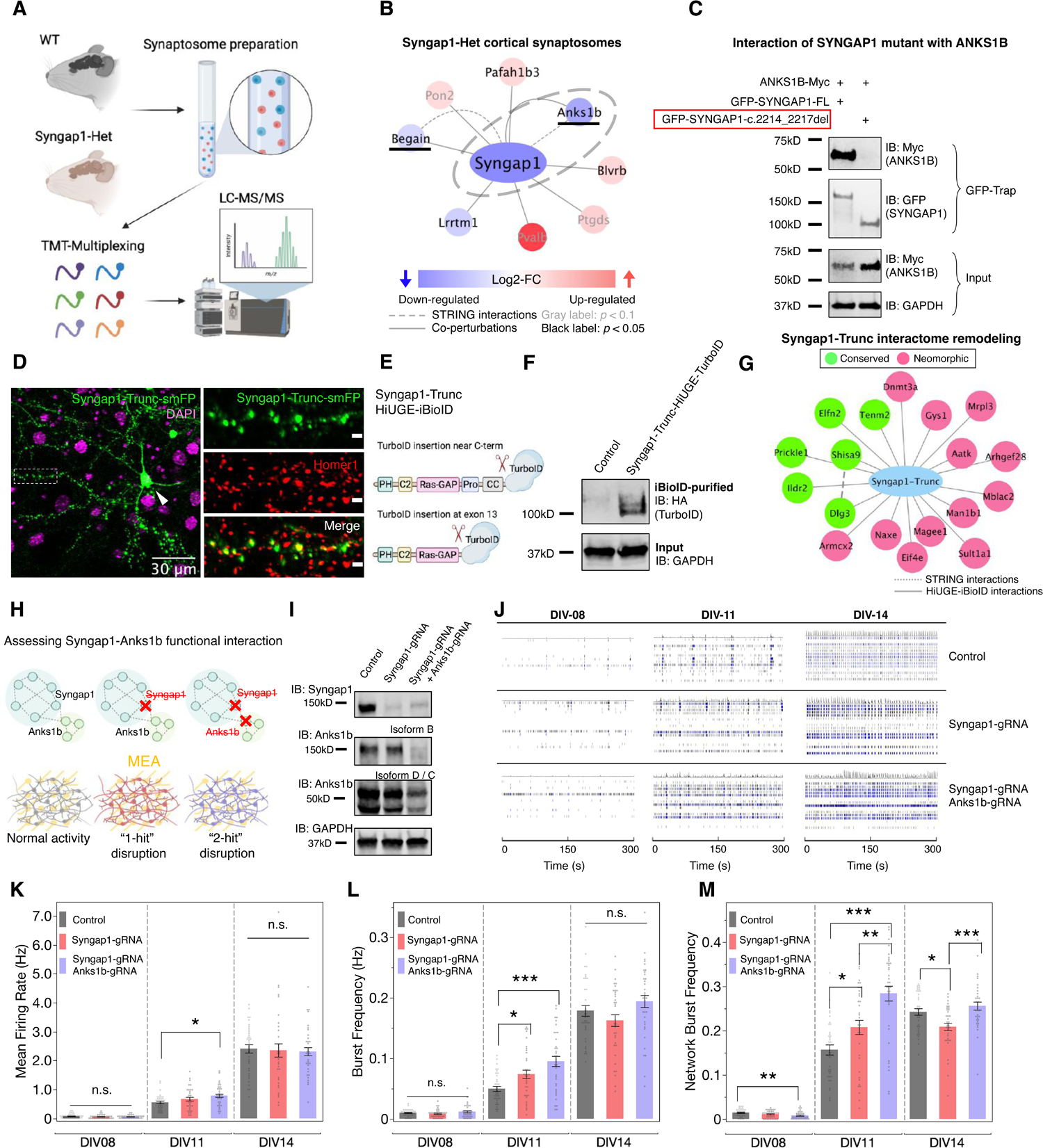
Proteomic co-perturbation and functional convergence of Syngap1 and Anks1b. (**A**) Schematic illustration of the quantitative proteomic characterization of Syngap1-Het synaptosomes. (**B)** Proteomic alterations identified in the Syngap1-Het synaptosome. Proteins that overlap with Syngap1 interactome are underlined. (**C**) Co-immunoprecipitation result showing loss of interaction with ANKS1B in frame-shifting c.2214_2217del SYNGAP1 mutation. (**D**) HiUGE labeling of truncated Syngap1 at exon 13 shows synaptic localization (boxed region) and aberrant somatic mis-localization (arrowhead). Scale bar in the enlarged view represents 2μm. (**E**) Schematic illustration of labeling truncated Syngap1 with TurboID by targeting exon 13. (**F**) Western blot showing TurboID-HA labeled Syngap1 truncation at the expected molecular mass. (**G**) Proximity interactome network showing conserved and neomorphic interactions in Syngap1 truncation. Note that interaction with Anks1b is no longer detected. (**H**) Schematics assessing phenotypes of the Syngap1-Anks1b functional interaction. (**I**) Western blot confirming disruption of Syngap1 and Anks1b expression. (**J**) Representative raster plots of neural activities at DIV-08, 11, and 14. (**K-M**) Neurometrics showing further depletion of Anks1b exacerbates the development of precocious neural activity associated with Syngap1-LOF. *: *p* < 0.05; **: *p* < 0.01; ***: *p* < 0.001; n.s.: non-significant. One-way ANOVA followed by *post-hoc* Tukey HSD tests (n = 36 wells). Plots are mean ± SEM.

As truncating mutations of Syngap1 confer significant genetic risk for autism ^50,55^, we next asked whether these mutations predicted to be pathogenic might lead to perturbation of its interactors, including Anks1b. When expressed in HEK293T cells, C-term truncation of human SYNGAP1 at amino acids (a.a.) 730 (C1-Trunc) and a.a. 848 (C2-Trunc) completely abolished binding with ANKS1B. In contrast, the interaction was retained with truncation at a.a. 1181 (C3-Trunc) or a truncation missing the N-term a.a. 2-361 (N-Trunc). These results suggest that the sequence surrounding the disordered region of SYNGAP1 (a.a. 848-1181) is crucial for the interaction with ANKS1B (Fig. S6). A pathological mutation found in a human patient (SYNGAP1-c.2214_2217del ^61^), which resulted in a frame-shift and premature truncation in this region, also abolished the SYNGAP1-ANKS1B interaction (Fig. 2C). A previous report showed that a 2 a.a. substitution in the C-term PDZ-binding motif of Syngap1 impaired several synaptic PPIs, but not Anks1b ^62^. As an orthogonal validation of the putative Anks1b interaction region identified above, we sought to test if a more disruptive truncation of Syngap1 that ablated this region could lead to a deeper remodeling of its interactome, including a loss of the Anks1b interaction *in vivo*. Endogenously expressed smFP-labeled Syngap1 truncated at exon 13 (Syngap1:1-744-smFP) retained synaptic localization in cultured neurons, although mis-localization in the soma was detected as well (Fig. 2D). We then targeted this locus to generate TurboID fusion to the truncated Syngap1 protein. HiUGE-iBioID revealed that although some interactors were preserved in Syngap1:1-744 (e.g., Dlg3, Shisa9, Prickle1), most of its synaptic interactions were abolished, including Anks1b (Fig. 2E-G, Table S1). Thus, we confirmed that the region identified by structure-function analysis *in vitro* is also crucial for the Syngap1-Anks1b interaction *in vivo*.

### Depletion of Anks1b exacerbates electrophysiological abnormalities associated with Syngap1 during *in vitro* neural development

We next sought to determine whether Syngap1 and Anks1b functionally interact by asking if further depletion of Anks1b would ameliorate or aggravate the phenotypes found in Syngap1 loss-of-function (LOF) mutants (Fig. 2H). AAV-mediated CRISPR disruption targeting Syngap1 at exon 13 and the first common exon of Anks1b (exon 15) led to a profound loss of their total protein (Fig. 2I). Using a multielectrode array (MEA) system to monitor electrophysiological activities longitudinally during neurodevelopment, we observed elevated firing rate and burst activities in Syngap1-gRNA treated neurons at DIV11, but not at DIV8 or DIV14 (Fig. 2J-M). This result phenocopied previous reports demonstrating that Syngap1 LOF atypically accelerated synaptic and network activities during the synaptogenesis period ^63,64^. Interestingly, further depletion of Anks1b exacerbated the abnormally heightened neural activity linked to Syngap1 mutation (Fig. 2J-M), primarily during the period of synaptogenesis (DIV 11). Taken together, the data support a model in which Syngap1 and Anks1b physically and functionally interact within a common pathway to regulate the development of neuronal activity. Based on the phenotype and common biological pathways found in the overlapping interactomes, this effect may be due to the altered developmental trajectory of the glutamate receptor module found in both networks.

### HiUGE-iBioID and spatial co-perturbation proteomics reveal targets that can restore spontaneous activity of neurons harboring a patient-derived Scn2a^+/R102Q^ mutation

Another critical rationale for interrogating endogenous interactomes is the prospect of identifying proteins functioning with the autism-implicated targets that can be modulated to buffer or normalize phenotypes. Such candidates could be relevant targets for future drug development. To test this possibility, we focused on Scn2a, mutations of which are one of the most highly significant for association with autism ^3,65–67^. In recent years, whole exome sequencing (WES) from autistic individuals identified a missense mutation, SCN2A-p.R102Q, that was previously reported ^68,69^ but uncharacterized. Clinical observations of one patient with this mutation included meeting DSM-5 diagnostic criteria for autism spectrum disorder (i.e., qualitative differences in social communication and the presence of restrictive interests/repetitive behaviors), minimal use of spoken language, disruptive and impulsive behaviors, sleep problems, and gastrointestinal concerns (reflux and constipation). A neurological exam including EEG did not reveal evidence of epilepsy. To test the mechanistic linkage of this mutation to autism, we generated a Scn2a point-mutant heterozygous mouse model (Scn2a*^+^*^/R102Q^) (Fig. 3A). Behavioral testing of the Scn2a^+/R102Q^ mice found that compared to WT littermates, the mutants presented with hyperactivity and reduced anxiety in the elevated zero maze, excessive repetitive behaviors in the self-grooming and the hole-board tests, and reduced ultrasonic vocalizations (USV) during social interactions (Fig. 3B). Social behavioral tests were also conducted with the resident-intruder and social dyadic assays where C3H/HeJ males served as social partners of WT and Scn2a^+/R102Q^ males. In both the resident-intruder and the dyadic tests, the social behaviors of the mutants appear to be relatively similar to WT, although abnormal numbers of social events and reduced withdrawal were noted (Fig. S7). Hence, the primary behavioral deficits of Scn2a^+/R102Q^ mice lie within social communication and repetitive behaviors. It is known that mutations in Scn2a can result in either infantile epileptic encephalopathy (IEE) or autism, depending upon the nature of the mutational effect (gain- or loss-of-function) ^70^. Cortical neurons cultured from Scn2a^+/R102Q^ mice exhibited a significant attenuation in neural firing activity (∼40% decrease at DIV14; *p* < 0.001, Fig. 3D), consistent with a role in autism but not IEE. Next, we analyzed the spatial proteome of these mutants versus WT mice using fractionation steps of whole brain tissue adapted from the Localisation of Organelle Proteins by Isotope Tagging after Differential ultraCentrifugation (LOPIT-DC) procedure ^30^, as we have published previously ^71^. Interestingly, we identified a down-regulation of a protein cluster associated with voltage-gated sodium channel (VGSC) activity in Scn2a^+/R102Q^ mice. This VGSC modulatory cluster included candidates intersecting with the Scn2a HiUGE-iBioID interactome: an auxiliary β subunit, Scn1b ^72^, and an intracellular VGSC modulator, Fgf12 ^73^ (Fig. 3C; Table S4).

**Fig. 3.**
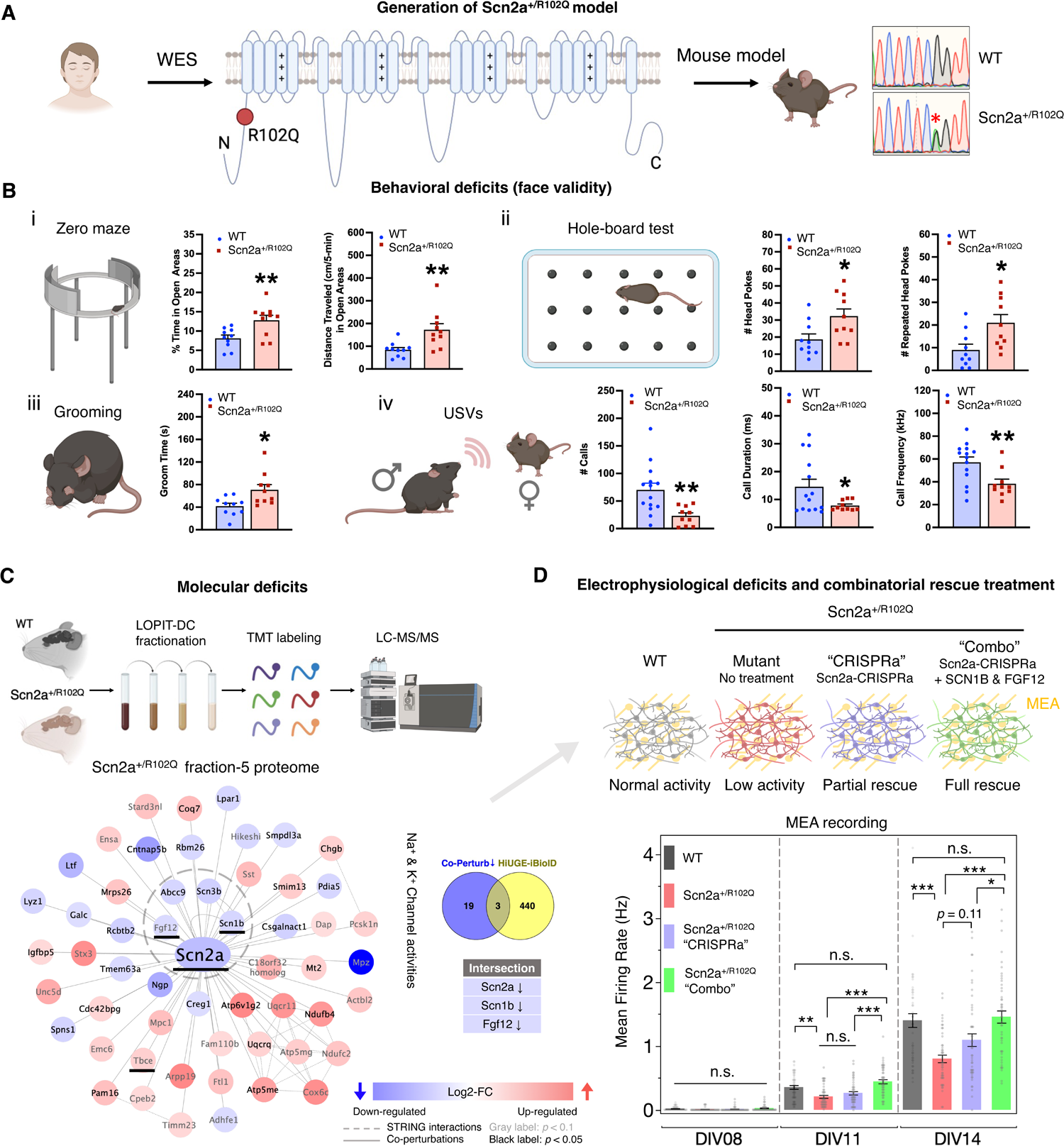
Intersectional proteomics reveal hidden molecular mechanism of a patient-derived Scn2a mutation. (**A**) Generation of a mouse model based on a clinically identified Scn2a missense mutation (R102Q) in autistic individuals. WES: whole exome sequencing. (**B**) Behavioral face-validity of Scn2a^+/R102Q^ mutants was assessed by (**i**) zero maze as the percent time and distance traveled in the open areas, (**ii)** hole-board test as numbers of head pokes and repeated head-pokes, (**iii)** self-grooming, and (**iv**) ultrasonic vocalizations (USVs) as numbers of calls, call durations, and call frequencies during social interaction. No differences were detected in the metrics of pre-social (baseline) responses in the USV tests. *: *p* < 0.05, **: *p* < 0.01, WT *vs.* Scn2a^+/R102Q^ mice; independent samples t-tests, two-tailed. Statistics are summarized in Table S6. (**C**) Spatial proteomics reveals co-perturbations in Scn2a^+/R102Q^ mutants. MCL analysis discovered a key cluster associated with voltage-gated channel activity that is down-regulated, including three targets that intersect with the Scn2a HiUGE-iBioID interactome (underlined). (**D**) Scn2a^+/R102Q^ mutant neurons show attenuated activity with the MEA. Scn2a-CRISPRa treatment and a “Combo” treatment with additional expression of SCN1B and FGF12 show differential efficacy in restoring neural activity deficits. *: *p* < 0.05; **: *p* < 0.01; ***: *p* < 0.001; n.s.: non-significant. One-way ANOVA followed by *post-hoc* Tukey HSD tests (n = 48 wells). Plots are mean ± SEM.

Based on the finding of the downregulated VGSC modulatory cluster, a rescue strategy to restore the electrophysiological deficits in Scn2a^+/R102Q^ neurons was devised. First, lentiviral-mediated CRISPR-activation (CRISPRa) was used to upregulate endogenous Scn2a expression (Fig. S8A). A similar strategy has recently been used to phenotypically rescue Scn2a haploinsufficiency ^74^; however, it has not yet been tested in neurons harboring patient-derived missense mutations. This treatment transcriptionally activates Scn2a expression; however, it is expected to amplify both the WT and mutant allele. RT-PCR data from WT and Scn2a^+/R102Q^ neurons showed normal transcript levels (Fig. S8B), suggesting the reduced protein levels in the mutant were likely due to post-transcriptional effects such as protein destabilization. Thus, it was unclear if Scn2a-CRISPRa alone would fully rescue phenotypes, as the effect potentially could be diluted by the mutant allele. Indeed, CRISPRa treatment only partially rescued the Scn2a^+/R102Q^ phenotype (Fig. 3D, purple bars), suggesting this approach is unlikely to work in the context of missense mutations. A treatment strategy was next tested by supplying either additional SCN1B or FGF12 via AAV-mediated expression to augment these down-regulated intersecting proteins identified in the VGSC modulatory cluster. MEA analysis revealed that neither expressing SCN1B or FGF12 alone, or when added together, was sufficient to rescue the phenotype (Fig S8C). Finally, a combinatory (“combo”) approach was tested, combining the upregulation of all three of the depleted VGSC cluster proteins. This combo approach resulted in a full restoration of neural firing activity metrics at DIV14 (Fig. 3D, S8D), suggesting the potential dominant-negative effect of the mutation in Scn2a-CRISPRa was overcome by the increased expression of SCN1B and FGF12; thereby confirming the hypothesis that this modulatory cluster is crucial for the phenotypic rescue of Scn2a^+/R102Q^. Collectively, these results strongly indicate that intersectional proteomics between HiUGE-iBioID and spatial co-perturbation can be an informative approach to discover novel approaches for the functional rescue of abnormal phenotypes and potential therapeutic targets.

## Discussion

Here we report a strategy combining the advantages of HiUGE and iBioID to resolve native interactomes associated with 14 autism-associated proteins. The combination of the interaction data presented for Syngap1 and Anks1b and the Scn2a rescue results validate the HiUGE-iBioID method for discovering the functional links between autism genetics and proteomics. The findings also emphasize an effective proteomic-driven systems-biology approach to discover molecular mechanisms and potential treatment targets.

Compared to immunoprecipitation or recombinant BioID expression methods *in vitro*, HiUGE-iBioID is expected to have four key benefits. First, the bait protein is expressed from the endogenous promoter with native cell-type specificity preserved at physiological levels. Second, the cells expressing the bait protein are within the context of the tissue, obviating perturbations to their native environment essential for development and cell physiology that can occur *in vitro*. Third, interactomes are covalently marked as they exist *in vivo* and thus, the proteins can be purified subsequently under stringent conditions without the need to optimize samples for weak or transient interactions. This feature is especially important for protein complexes organized by transmembrane baits, which must be extracted from the lipid bi-layer under conditions that are unfavorable for maintaining many PPIs. Fourth, the selection of the bait-protein is not limited by viral packaging capacity or availability of high-specificity and validated antibodies.

HiUGE-iBioID is a technique that, from a biochemical perspective, combines the strengths of endogenous protein-based purification from tissue that antibodies afford with the advantages of covalent marking of proximity interactomes by BioID. These qualities suggest that HiUGE-iBioID may be an optimal approach. Nevertheless, an essential question is how its interactomes compare with analogous studies using immunoprecipitation or recombinant BioID expression. To address this important issue, we performed a direct comparison of HiUGE-iBioID to antibody immunoprecipitations of Anks1b and Scn2a from mouse brain tissue (Fig. S4 I, J). While there was a significant overlap in the interactomes between HiUGE-iBioID and each immunoaffinity pulldown, the differences between them were even greater (Fig. S4 K, L). For example, immunoprecipitation of the voltage-gated sodium channel, Scn2a, identified the expected proteins that overlap with HiUGE-iBioID, including clusters of Na^+^ / K^+^ channels and Fgf12, consistent with previous reports ^28,75^. However, GO analysis of the fraction that did not overlap with HiUGE-iBioID yielded top-ranked molecular function terms of “structural constituent of the ribosome” and multiple RNA processes. In contrast, the fraction unique to HiUGE-iBioID were enriched correctly for top GO terms of voltage-gated ion channel functions. Similar conclusions were noted from the comparison of HiUGE-iBioID to the recent publication of immunoprecipitation-based interactomes of Syngap1, Shank3, Scn2a, and Ctnnb1 ^28^ (Fig. S9). We also compared recently published data of recombinant bait-BioID expression in cultured mouse neurons to HiUGE-iBioID for the baits Syngap1, Shank3, and Lrrc4c that were shared between both studies ^27^ (Fig. S10). While the BioID and HiUGE-iBioID interactomes had more in common with each other than the analogous immunoprecipitation comparisons, there were again more differences between the interactomes than there were similarities. While HiUGE-iBioID specific interactomes for each had the top GO term “Glutamate receptor binding” for all three synaptic baits, the recombinant BioID specific data were enriched for top GO terms such as “Tubulin binding”, “RNA binding”, “Heat shock protein binding”, and “Acidic amino acid transmembrane transporter activity”. While further comparisons are needed to confirm the above observations, the available data suggest that endogenous proximity proteomics outperforms both immunoprecipitation and recombinant BioID methods *in vitro*. Employing HiUGE-iBioID for the *in vivo* study of endogenous protein complexes appears to be an advantageous method overall.

Consistent with the observation that HiUGE-iBioID performs well, each interactome revealed the expected, as well as new potential biological insights following further analysis with MCL to partition biologically-relevant communities (Fig. S11-24). For example, neurobeachin (Nbea) (Fig. S16) is known to regulate the surface levels of ionotropic GABA and glutamate receptors as well serve as an A-Kinase (PKA) Anchoring Protein (AKAP) ^76–79^. HiUGE-iBioID revealed that both the regulatory and catalytic subunits of PKA were detected. Clusters of the relevant receptors with significant enrichment for “GABA signaling pathway”, “Ionotropic glutamate receptor signaling”, and “Regulation of AMPA receptor activity” were also found, consistent with Nbea’s known functions. Of note, the “GABA signaling pathway” cluster was also significantly enriched for the terms “autism spectrum disorder” and “epilepsy”, both to which Nbea is implicated ^80,81^. Interestingly, GABA and glutamate surface levels are thought to be modulated by distinct pathways influenced by Nbea ^82^. Consistent with this idea, clusters enriched for “AP-type membrane coat adaptor complex”, “COPI vesicle coat”, and “TRAPP complex” were discovered, suggesting these may be the trafficking processes that Nbea modulates. Moreover, the analysis suggests Nbea may influence other signaling pathways that have yet to be appreciated, including “G-protein-coupled GABA receptors” and “Voltage-gated potassium channel complexes”, although additional experimental analyses will be needed to test these new interactome-derived hypotheses.

In addition to specificity advantages, HiUGE-iBioID is easily scalable in terms of time and resources when compared to the alternative of the traditional transgenic animal approaches to generate endogenous protein fusions *in vivo*. We have found that forebrain tissue collected from as few as two mice is sufficient for one biological replicate. Therefore, a typical mouse litter (∼ 8 pups) is sufficient to analyze a bait interactome in biological triplicates. We anticipate this method can be optimized even further for larger-scale applications, where costs can be limiting. These approaches could include multiplexed mass-spectrometry techniques such as isobaric labeling ^83,84^, or new advancements in mass spectrometry scan rates that reduce instrument time.

We also recognize a few limitations of our method. First, the proximity proteomes reported here are based on an enrichment with the bait over negative control, thus it is possible some proteins are present yet not identified as significantly enriched ^85^. Second, since the fusion proteins are expressed at an endogenous level, efficient biotinylation may be challenging for some low-expression proteins. Third, common to all protein fusion strategies, it is unrealistic to guarantee that the tag does not alter any interaction. The ability to generate fusion proteins by inserting TurboID directly into coding exons, or by splicing TurboID in from introns for proteins that cannot be altered at their N- or C-termini, provides flexible gRNA selection as well as protein engineering design. We envision further developments to our approach will include adapting *in silico* structure prediction tools (e.g. AlphaFold ^86^ and ESMFold ^87^) to minimize potential disturbances by structure-based optimization of TurboID fusions. Additionally, rapid advancements in the sensitivity of mass-spectrometry ^88–90^ promise to enable further HiUGE-iBioID analysis, even for low-level proteins.

A crucial finding from the current HiUGE-iBioID application is that diverse autism genes physically form protein interaction networks with each other and can be co-perturbed in genetic models of autism. This result confirms that divergent genetic mutations can converge at the protein level to drive autism neurobiology, providing a working paradigm to prioritize co-regulated candidates in physiology and atypical brain development. Furthermore, we demonstrate that high-confidence autism genes interact with other lower-confidence autism genes at the proteome level. Indeed, three proteins detected in the interactome dataset (Itsn1, associated with Scn2a, Scn8a, Ank3, and Shank3 baits; Nav3, associated with Scn2a bait; and Hnrnpul2, associated with Hnrnpu bait) were discovered subsequently as new autism risk genes of genome-wide significance ^38^ during the preparation of this manuscript. These results emphasize the possibility that additional genes - whose significance in autism are as yet unknown - exist in the dataset. Thus, HiUGE-iBioID may be useful to prioritize lower-confidence autism genes, which could either play a role in regulating core autism drivers or serve as novel targets for pharmacological developments. In addition, we expect the endogenous autism interactome data will stimulate a new impetus for predictive modeling of autism genetics ^91,92^ and cell-type specific analyses harnessing single-cell proteomics ^93–95^.

Informed by the autism interactome and proteomic co-perturbation results, we focused on a tightly co-regulated pair, Syngap1 and Anks1b ^27,46–48^, and verified the functional significance of their interaction. We identified a putative internal region on Syngap1 that is crucial for interaction with Anks1b, and confirmed that Syngap1 truncation led to remodeling of its interactome, including a loss of interaction with Anks1b. The data support the notion that remodeling of synaptic protein complexes, such as previously reported PDZ-associated alterations ^62^ and the disruption of specific interactions like Syngap1-Anks1b, could be possible mechanisms relevant to Syngap1 synaptopathy beyond nonsense-mediated decay ^50^. Further, we identified several neomorphic proteomic interactions associated with the Syngap1 truncation (Fig. 2G). The mechanisms of whether or how these abnormal interactions might contribute to Syngap1-associated phenotypes remain to be carefully assessed, and cannot be easily extrapolated to patients. The additive effects of Syngap1 and Anks1b deficits seen in the aberrant neural activity indicate that the downregulation of Anks1b in Syngap1-Het mice is not a compensatory effect, but rather it may contribute to a mechanism that alters neuronal development. The disordered region of Syngap1, identified as a critical domain mediating binding with Anks1b, contains proline-rich stretches, a poly-histidine motif, and is phosphorylated at multiple sites ^96,97^. This architecture suggests their interaction could be under activity-dependent kinase modulation. Accordingly, within the Anks1b-Syngap1 core interactome, there are 12 kinases / phosphatases, 4 of which are likely autism risk genes themselves (Pak2, Cdkl5, Ntrk3, Dyrk1a) ^2^. Thus, an intriguing speculation is that these gene products may play a role in regulating synaptogenesis via acting on the Syngap1/Anks1b complex, and contributing to the activity-dependent shuttling of Syngap1/Anks1b during plasticity.

A notable finding of this study is the discovery of a mechanism for restoring *in vitro* neural activity deficits associated with a patient-derived SCN2A-p.R102Q mutation. We have identified three proteins (Scn2a, Scn1b, and Fgf12) within the VGSC complex that are critical for phenotypic rescue by intersecting the Scn2a HiUGE-iBioID interactome with co-perturbation spatial-proteomics in a mouse model harboring this mutation. A qPCR analysis revealed that the abundances of Scn2a, Scn1b, and Fgf12 mRNAs in cultured Scn2a^+/R102Q^ neurons are comparable to the WT (Fig. S8B), suggesting the downregulation of these proteins occurs at the post-transcriptional level - likely due to destabilization of VGSC supramolecular assembly. The exact mechanisms as to how the loss of a positively charged arginine residue on the N-term intracellular tail of Scn2a affects VGSC assembly remains to be determined. Similarly, the effects of these perturbations need to be assessed further as some alterations may be benign with respect to phenotypes.

Both β-subunits and FGF family proteins play critical roles in maintaining neuronal excitability through regulating VGSC kinetics ^72,98^. Thus, their downregulation in Scn2a^+/R102Q^ is likely a contributing factor to VGSC channelopathy. These results also explain why Scn2a-CRISPRa treatment alone is insufficient to rescue the electrophysiological deficits, especially since the treatment does not differentiate the functional allele from the point-mutant allele ^74^. We further show that supplying additional SCN1B and FGF12 without Scn2a-CRISPRa does not rescue the deficits either (Fig. S8C). Thus, it appears that upregulation of all three components is required to provide the molecular environment necessary to restore functional VGSC complexes and reinstate the level of spontaneous neural activity. The *in vitro* phenotypic rescue presented here will require future testing *in vivo*. Although we did not observe an overt epileptogenic effect on the MEA, these potential adverse effects should be carefully assessed in future animal studies. In addition, since VGSCs are believed to contribute to neuronal excitability and plasticity at both pre- and post-synapses ^65,99^, future studies are needed to further dissect their unique subcellular effects.

Together, our results show that HiUGE-iBioID provides a new CRISPR/proximity proteomics method to reveal native proteomes *in vivo* with higher-confidence interactomes than immunoprecipitation or BioID over-expression approaches, and with considerable ease compared to traditional transgenic approaches. Combined with co-perturbation proteomics, our intersectional approach offers a generalizable strategy to identify and prioritize candidates for discovering new biology and potential therapeutic targets. Thus, future work could adopt a similar approach to investigate genetic co-perturbations and PPIs in other models. We expect that the framework developed in this study will encourage further research of native proximity interactomes in the brain, enhancing our understanding of proteome organization in various aspects of cellular neurobiology and disease.

## Acknowledgments

We would like to thank Drs. Tyler Bradshaw and Jamie Courtland for performing synaptosome and fractionation proteomics, Dr. Ramona M. Rodriguiz, Mr. Christopher R. Means, Mr. Nathan Franklin, and Ms. Sarah Page Steffens for conducting the behavioral experiments, and Dr. Rodriguiz for helping with the behavioral statistical analyses. We would like to thank Arinze Okafor for assistance with the similarity matrix plot, Greg Waitt for developing the DPMSR Proteome Discoverer Data Visualization Tool, and Daniel J. Morone for assistance with lentiviral production and titration. We would also like to thank Dr. Jiechun Zhou for her assistance with mouse husbandry. Schematic illustrations were created using a licensed BioRender.com account.

## Data and Materials Availability

Requests for data, resources, and reagents should be directed to and will be fulfilled by the Corresponding Author, Dr. Scott Soderling. Key plasmids from this study have also been deposited to Addgene.

## Code availability

Requests for custom computer code used in this study should be directed to and will be fulfilled by the Corresponding Author, Dr. Scott Soderling. Key codes from this study have also been deposited to Github.

## Author contributions

YG and SHS conceived the study. YG, MT, DS and EJS performed HiUGE-iBioID experiments and analyses. SAGM designed CRISPRa constructs and JZ performed validation experiments. YG and MT performed MEA experiments. WCW designed and managed the behavioral experiments and statistical analyses. GD and YJ contributed to the generation of Scn2a^+/R102Q^ model. DV and LGS performed gene-set analyses. CAG, JDB, and SHS supervised the collaborative study. YG and SHS wrote the original manuscript. All authors contributed to manuscript editing and revision.

## Materials and Methods

### Animals

For all CRISPR-Cas9-related experiments, H11-Cas9 mice (Jackson Laboratory #28239) were used. The Syngap1-Het mouse model, originally described by Kim and colleagues ^56^, was a gift from Dr. Gavin Rumbaugh. The Scn2a^+/R102Q^ mouse model of the human c.305G>A (p.R102Q) mutation ^68,69^ was created by the Duke Transgenic and Knockout Mouse Shared Resource. The Scn2a^+/R102Q^ mice were generated using a heterozygous breeding scheme and genotyping was performed by sequencing the amplicon using the following primer set: Scn2a-s, acagacatggcggaaaacatgag; and Scn2a-as, agcaggagaggaaagaaagaagc. C3H/HeJ males (#000659; Jackson Laboratory, Bar Harbor, ME) served as social partners in the resident-intruder and social dyadic tests. All procedures were performed with a protocol approved by the Duke University Institutional Animal Care and Use Committee in accordance with US National Institutes of Health guidelines.

### Single-vector HiUGE TurboID knock-in

HiUGE plasmids were constructed based on the previously described method ^31^. Briefly, a donor of HA-tagged TurboID coding sequence was flanked by DNA sequences that were specifically recognized by a synthetic donor-specific gRNA (DS-gRNA), inert to the host genome. The gene-specific gRNA (GS-gRNA) expression cassette (U6 promotor, GS-gRNA, and gRNA scaffold) was inserted in tandem to the DS-gRNA expression cassette permitting a single-vector delivery. GS-gRNAs were designed using CRISPOR ^100^, and a pair of 23-24mer oligonucleotides were annealed and ligated into the *SapI* site of the GS-gRNA expression cassette. The genomic target sequences for the baits were: Anks1b: cggcgggtatcagaaaatcgtgg, Shank2: aacagctgctggacagataaggg, Shank3: cgtgcgctcaggcagctggatgg, Nckap1: gcatttctcagcaacacataagg, Nbea: ggcattatgagcatcagaacagg, Arhgef9: cattctggcaaaacttcagtagg, Ank3: gaagaaggaaatccggaacgtgg, Scn2a: ggacaaggggaaagatatcaggg, Scn8a: tcagggagtccaagtgctagagg, Hnrnpu: tggagtcagcattatcaccaagg, Homer1: ttagctgcattccagtagcttgg, and Wasf1: gttcgatgaagtagactggctgg. To protect the PDZ-binding motifs of Syngap1, Ctnnb1, and Iqsec2, an intron-targeting strategy ^33^ was used, where the donor was flanked by intron / exon boundary sequences from an obligatory intron to enable internal TurboID insertion. The following intronic sequences were targeted: Syngap1: acttattgagacgcttcgcgggg, Ctnnb1: aacaggcttccagatgcgatggg, and Iqsec2: agggccaactccaaatagggagg. To insert TurboID while protecting the C-term PDZ-binding motif of Lrrc4c that has only one coding exon, a custom donor was used where the coding sequence of a.a. ETQI was appended to the C-term of the TurboID donor, thus preserving the native motif. The genomic target for Lrrc4c: gagttcattcggatcaataacgg. Exon 13 of Syngap1 was targeted for Syngap1 truncation and disruption: acggactcggtctcagcccatgg. To endogenously express soluble TurboID as a survey for background detection, C-term or 3’-UTR sequences were targeted with a stop codon - internal ribosome entry site (IRES) - TurboID-HA donor. The genomic target sequences were: Syngap1: aggaggtctgtgacgctgggtgg, Scn2a: agtttggcatagacctcctgagg, Hnrnpu: aaacagtcgacttcttgtgaagg, Ctnnb1: ttataagctttcttacctaaagg, Iqsec2: tactggggagcaggatagtctgg, Homer1: ttagctgcattccagtagcttgg, and Wasf1: gttcgatgaagtagactggctgg.

### AAV preparation

AAV was prepared following previously described methods ^23,31^. Briefly, concentrated AAV virus was produced in HEK293T cells grown in six 15cm dishes by triple-transfection with 15μg HiUGE vector, 30μg pADdeltaF6, and 15μg serotype plasmid (pUCmini-iCAP-PHP.eB, a gift from Viviana Gradinaru ^101^, Addgene plasmid #103005). Three days following transfection, cells were lysed and virus was concentrated using an Optiprep density gradient (Sigma #D1556). Small-scale AAV virus was produced in HEK293T cells grown in 12-well plates by triple-transfection with 0.4μg HiUGE vector, 0.8μg pADdeltaF6, and 0.4μg serotype 2/1 plasmids. Three days following transfection, the virus-containing medium was filtered through Costar Spin-X columns (Sigma #8162) as previously described ^31^.

### AAV injection and biotin injection for HiUGE-iBioID

Neonatal (P0-2) H11-Cas9 mice were injected intracranially with the purified AAV (2µL per hemisphere, PHP.eB serotype, > 10^10^ GC / μL titer). Donor vector backbone AAV (empty gRNA) was used as negative control. Approximately 3 weeks after injection, mice received daily intraperitoneal injections (i.p.) of biotin (50mg/kg) over 5 consecutive days. Forebrain tissue was collected one day after the final injection and snap-frozen at −80°C until purification.

### HiUGE-iBioID sample purification

For each replicate, forebrain tissue from two mice was combined, homogenized, sonicated in RIPA lysis buffer (Cell Signaling #9806) supplemented with cOmplete protease inhibitor cocktail (Sigma #11873580001), and centrifuged at 16,000 xg for 30min at 4°C. The supernatant lysate was desalted with Zebra columns 7K MWCO (ThermoFisher #89894 or #89892). The flow-through was combined with 150µL magnetic Strepavidin beads (Pierce #88816) and incubated at 4°C overnight. On the next day, the beads were washed with the following steps: RIPA buffer 2 times, 1M KCl once, 0.1M Na_2_CO_3_ once, 2M urea in 10mM Tris-HCl once, and RIPA buffer 2 times. Biotinylated proteins were eluted by boiling the beads in 90µL 2X sample buffer, supplemented with 2.5mM biotin, and used for downstream LC-MS/MS and Western blot analyses.

### HiUGE-iBioID LC-MS/MS analysis

Samples were spiked with undigested bovine casein at a total of either 120 or 240 fmol as an internal quality control standard. Next, samples were supplemented with 12.4 μL of 20% SDS, reduced with 10 mM dithiolthreitol for 30 min at 80 °C, alkylated with 20 mM iodoacetamide for 30 min at room temperature (RT), then supplemented with a final concentration of 1.2% phosphoric acid and 723 μL of S-Trap (Protifi) binding buffer (90% MeOH/100mM TEAB). Proteins were trapped on the S-Trap micro cartridge, digested using 20 ng/μL sequencing grade trypsin (Promega) for 1 hr at 47 °C, and eluted using 50 mM TEAB, followed by 0.2% FA, and lastly using 50% acetonitrile (ACN) /0.2% FA. All samples were lyophilized to dryness. Samples were resolubilized using 12 μL of 1% TFA/2% ACN with 25 fmol/μL yeast ADH.

Quantitative LC-MS/MS was performed on 2 μL (∼17% of total sample) using a nanoAcquity UPLC system (Waters Corp) coupled to a Thermo Orbitrap Fusion Lumos high resolution accurate mass tandem mass spectrometer (Thermo) equipped with a FAIMSPro device via a nanoelectrospray ionization source. Briefly, the sample was first trapped on a Symmetry C18 20 mm × 180 μm trapping column (5 μl/min at 99.9/0.1 v/v water/ACN), after which the analytical separation was performed using a 1.8 μm Acquity HSS T3 C18 75 μm × 250 mm column (Waters Corp.) with a 90-min linear gradient of 5 to 30% ACN with 0.1% formic acid at a flow rate of 400 nanoliters/minute (nL/min) with a column temperature of 55°C. Data collection on the Fusion Lumos mass spectrometer was performed for three difference compensation voltages (−40v, −60v, −80v). Within each CV, a data-dependent acquisition (DDA) mode of acquisition with a r=120,000 (@ m/z 200) full MS scan from m/z 375 – 1600 with a target AGC value of 4e5 ions was performed. MS/MS scans were acquired in the ion trap in rapid mode with a target AGC value of 1e4 and max fill time of 35 msec. The total cycle time for each CV was 0.66s, with total cycle times of 2 sec between like full MS scans. A 20s dynamic exclusion was employed to increase depth of coverage. The total analysis cycle time for each injection was approximately 2 hours.

Following UPLC-MS/MS analyses, data were imported into Proteome Discoverer 2.5 (“PD”, Thermo Scientific Inc.) and individual LCMS data files were aligned based on the accurate mass and retention time of detected precursor ions (“features”) using Minora Feature Detector algorithm in Proteome Discoverer. Relative peptide abundance was measured based on peak intensities of selected ion chromatograms of the aligned features across all runs. The MS/MS data was searched against the SwissProt *M. musculus* database and a common contaminant/spiked protein database (bovine albumin, bovine casein, yeast ADH, etc.), and an equal number of reversed-sequence “decoys” for false discovery rate determination. Sequest with Infernys enabled (v 2.5, Thermo PD) was utilized to produce fragment ion spectra and to perform the database searches. Database search parameters included fixed modification on Cys (carbamidomethyl) and variable modification on Met (oxidation). Search tolerances were 2ppm precursor and 0.8Da product ion with full trypsin enzyme rules. Peptide Validator and Protein FDR Validator nodes in Proteome Discoverer were used to annotate the data at a maximum 1% protein false discovery rate based on q-value calculations. Note that peptide homology was addressed using razor rules in which a peptide matched to multiple different proteins was exclusively assigned to the protein has more identified peptides. Protein homology was addressed by grouping proteins that had the same set of peptides to account for their identification. A master protein within a group was assigned based on % coverage.

Prior to imputation, a filter was applied such that a peptide was removed if it was not measured in at least 2 unique samples (50% of a single group). After filtration, any missing data missing values were imputed using the following rules; 1) if only a single signal was missing within the group of three, an average of the other two values was used or 2) if two out of three signals were missing within the group of three, a randomized intensity within the bottom 2% of the detectable signals was used. To summarize to the protein level, all peptides belonging to the same protein were summed into a single intensity. This protein value was then subjected to a robust mean normalization in which the top and bottom 10 of the signals were removed and then the remaining mean was made to be the same across all samples.

The results were log2-transformed and analyzed using the PolySTest online tool ^102^. Proteomic detection was defined as proteins identified by at least 2 peptides in LC-MS/MS. Protein abundances were considered significantly enriched if they met FDR < 0.05 (PolySTest), and fold change ≥ 2 (Log2-FC ≥ 1) compared to controls. The enriched gene lists were filtered against known experimental and omnipresent biological contaminants, and soluble TurboID background detections. Note that this high-stringency filter removed a few well-known interactors such as Dlg4, Shank2, and Shank3 from the dataset of several baits.

### Synaptosomal preparation and proteomic analysis of Syngap1-Het mice

Synaptosomal preparation was performed with four adult Syngap1-Het mice and four WT controls. Briefly, rapidly isolated brain tissue was sliced to 1mm sections using a brain matrix (Zivic Instruments), followed by dissection of cortical and striatal tissue. Tissue was homogenized in homogenization buffer (320 mM sucrose, 5 mM HEPES, pH 7.4) using a Dounce homogenizer. The lysate was centrifuged at 1,000 xg to remove cell debris and nuclei. The supernatant was further centrifuged at 12,000 xg to obtain a crude synaptosomal pellet and it was resuspended in Tris buffer (320 mM sucrose, 5 mM Tris/HCl, pH 8.1). Additional centrifugation in a sucrose density gradient (0.8/1.0/1.2 M) at 85,000 xg was performed to isolate purified synaptosome at the 1.0/1.2 interface. The purified synaptosomes were subjected to multiplexed LC-MS/MS quantification following tandem mass tags (TMT) isobaric labeling.

### LOPIT-DC subcellular fractionation and proteomic analysis of Scn2a^+/R102Q^ mice

LOPIT-DC fractionation was performed with three adult Scn2a^+/R102Q^ mice and three WT controls, following a previously described method ^30,71,103^. Briefly, rapidly isolated brain tissue was added to isotonic TEVP homogenization buffer (320 mM sucrose, 10 mM Tris, 1 mM EDTA, 1 mM EGTA, 5 mM NaF, pH7.4), supplemented with cOmplete protease inhibitor cocktail (Sigma #11873580001). The tissue was homogenized for 15 passes in a 2mL Dounce homogenizer. The volume of the homogenate was brought up to a 5 mL volume with TEVP buffer, and then passed through a 0.5 mL ball-bearing homogenizer for two passes (14 µm ball, Isobiotec). Differential centrifugation steps were performed at 4°C sequentially at 200, 1000, 3000, 5000, 9000, 12,000, 15,000, 30,000, 79,000, and 120,000 xg. Fraction-5, determined to be enriched with Scn2a in a pilot experiment ^71^, was used for tandem mass tag (TMT)-multiplexed proteomic analysis.

### TMT-multiplexed quantitative LC-MS/MS analysis

Samples were supplemented with 100 μL of 8 M urea and probe sonicated. Protein concentrations were determined via Bradford Assay and ranged from 1.8 – 2.3 mg/mL. Samples were normalized to 120 μg using 8 M urea and spiked with undigested bovine casein at a total of either 120 or 240 fmol as an internal quality control standard. Next, they were supplemented with 13 μL of 20% SDS, reduced with 10 mM dithiolthreitol for 45 min at 32 °C, alkylated with 20 mM iodoacetamide for 45 min at RT, then supplemented with a final concentration of 1.2% phosphoric acid and 70 μL of S-Trap (Protifi) binding buffer (90% MeOH/100mM TEAB). Proteins were trapped on the S-Trap micro cartridge, digested using 100 ng/μL sequencing grade trypsin (Promega) for 1 hr at 47 °C, and eluted using 50 mM TEAB, followed by 0.2% FA, and lastly using 50% CAN/0.2% FA. All samples were then lyophilized to dryness.

For TMT labeling, each sample was resuspended in 120 μL 200 mM triethylammonium bicarbonate, pH 8.0 (TEAB). 40uL of each sample was combined to form an SPQC pool, which was then aliquoted into 3 SPQC pools. Fresh TMT10plex reagents (0.8 mg for each 10-plex reagent) were resuspended in 41 μL 100% ACN and was added to 75 μg of each sample. Samples were incubated for 1 hour at RT. After 1-hour reaction, 8 μL of 5% hydroxylamine was added and incubated for 15 minutes at RT to quench the reaction. Samples were combined then lyophilized to dryness.

For offline fractionation, samples were resuspended in 300uL 0.1% formic acid. 400μg was fractionated into 48 unique high pH reversed-phase fractions using pH 9.0 20 mM ammonium formate as mobile phase A and neat ACN as mobile phase B. The column used was a 2.1 mm x 50 mm BEH C18 (Waters) and fractionation was performed on an Agilent 1100 HPLC with G1364C fraction collector. Throughout the method, the flow rate was 0.4 mL/min and the column temperature was 55 °C. The gradient method was set as follows: 0 min, 3%B; 1 min, 7% B; 50 min, 50%B; 51 min, 90% B; 55 min, 90% B; 56 min, 3% B; 70 min, 3% B. Forty-eight fractions were collected in equal time segments from 0 to 52 minutes, then concatenated into 12 unique samples using every 12th fraction. For instance, fraction 1, 13, 25, and 37 were combined, fraction 2, 14, 26, and 38 were combined, etc. Fractions were frozen and lyophilized overnight. Samples were resuspended in 50 μL 1%TFA/2% ACN prior to LC-MS analysis.

Quantitative LC-MS/MS was performed on 2 μL (1 μg) of each sample, using a nanoAcquity UPLC system (Waters Corp) coupled to a Thermo Orbitrap Fusion Lumos high resolution accurate mass tandem mass spectrometer (Thermo) equipped with a FAIMSPro device via a nanoelectrospray ionization source. Briefly, the sample was first trapped on a Symmetry C18 20 mm × 180 μm trapping column (5 μl/min at 99.9/0.1 v/v water/ACN), after which the analytical separation was performed using a 1.8 μm Acquity HSS T3 C18 75 μm × 250 mm column (Waters Corp.) with a 90-min linear gradient of 5 to 30% ACN with 0.1% formic acid at a flow rate of 400 nanoliters/minute (nL/min) with a column temperature of 55°C. Data collection on the Fusion Lumos mass spectrometer was performed for three difference compensation voltages (−40v, −60v, −80v). Within each CV, a data-dependent acquisition (DDA) mode of acquisition with a r=120,000 (@ m/z 200) full MS scan from m/z 375 – 1600 with a target AGC value of 4e5 ions was performed. MS/MS scans were acquired in the Orbitrap at r=50,000 (@ m/z 200) from m/z 100 with a target AGC value of 1e5 and max fill time of 105 msec. The total cycle time for each CV was 1s, with total cycle times of 3 sec between like full MS scans. A 45 sec dynamic exclusion was employed to increase depth of coverage. The total analysis cycle time for each sample injection was approximately 2 hr.

Data were imported into Proteome Discoverer 2.4 (Thermo Scientific Inc.) and individual LCMS data files were aligned based on the accurate mass and retention time of detected precursor ions (“features”) using Minora Feature Detector algorithm in Proteome Discoverer. Relative peptide abundance was measured based on peak intensities of selected ion chromatograms of the aligned features across all runs. The MS/MS data was searched against the SwissProt M. musculus database (downloaded in Nov 2019), a common contaminant/spiked protein database (bovine albumin, bovine casein, yeast ADH, etc.), and an equal number of reversed-sequence “decoys” for false discovery rate determination. Mascot Distiller and Mascot Server (v 2.5, Matrix Sciences) were utilized to produce fragment ion spectra and to perform the database searches. Database search parameters included fixed modification on Cys (carbamidomethyl) and variable modification on Met (oxidation), Asn/Gln (deamindation), Lys (TMT6plex) and peptide N-termini (TMT6plex). Peptide Validator and Protein FDR Validator nodes in Proteome Discoverer were used to annotate the data at a maximum 1% protein false discovery rate based on q-value calculations. Note that peptide homology was addressed using razor rules in which a peptide matched to multiple different proteins was exclusively assigned to the protein has more identified peptides. Protein homology was addressed by grouping proteins that had the same set of peptides to account for their identification. A master protein within a group was assigned based on percent coverage. To account for any missing data (from a misalignment, low quality peak, low signal to noise, etc.) missing values were imputed by using a randomized intensity within the bottom 2% of the detectable signals. The data was then intensity normalized using a trim-mean normalization in which the highest and lowest 10% of the signals from each sample was excluded and then the remaining average intensities of the proteins was made equal across all of the samples.

The results were then analyzed using the Duke Proteomics and Metabolomics Shared Resource (DPMSR) Proteome Discoverer Data Visualization Tool. Proteomic detection was defined as proteins identified by at least 2 peptides in LC-MS/MS following a 1% FDR correction. Protein abundance was considered significantly altered if they met *p*-value < 0.05 (two-tailed *t*-test), and abs(fold change) ≥ 1.2 (abs(Log2-FC) ≥ 0.263). Those with *p*-value < 0.1 and abs(fold change) ≥ 1.2 (abs(Log2-FC) ≥ 0.263) were considered indicative candidates.

### Protein network visualization

Protein networks were constructed using Cytoscape ^104^ with nodes representing enriched or dysregulated proteins identified by LC-MS/MS. Known interactions with high confidence (i.e., 0.7 score) between these nodes were queried from the full STRING database (https://string-db.org) ^34,35^ and plotted on the network figure. To assess the percentage of interactions not reported in STRING queries, a low confidence (0.15) threshold was used. The Markov Cluster Algorithm in STRING and gene ontology (GO) analysis were used to detect protein communities within each interactome, with additional adjustments made based on known protein functions.

### Gene set enrichment analyses

Gene ontology (GO) enrichment of interactomes was searched against a custom statistical domain of all identified brain proteins (9,686 unique proteins, Table S5) from cumulative mouse brain proteomic studies in our lab (n=107), using ShinyGO ^105^. Pathway size boundary was set at between 10 and 500 to exclude ambiguous terms for querying the Molecular Function pathway database. Default pathway size boundary was used for all other queries. Redundancy removal option was enabled. SynGO function analyses ^106^ were also conducted with the synaptic bait proteomes versus the nucleus Hnrnpu proteome as control. Overlaps of identified interactors with the SFARI gene list (2022 Q1 release) were calculated using Venny (Oliveros, J.C. (2007-2015) Venny. An interactive tool for comparing lists with Venn’s diagrams. https://bioinfogp.cnb.csic.es/tools/venny/index.html). Hypergeometric tests were performed using an online tool (http://www.nemates.org/MA/progs/overlap_stats.html). In addition, we mapped autism risk genes identified at FDR < 0.1 (102 genes) in Satterstrom et al., 2020 ^3^ and FDR < 0.05 (185 genes) in Fu et al., 2022 ^7^ to mouse orthologs, and examined their overlap with each set of proteins obtained from the 14 HiUGE-iBioID experiments, via hypergeometric testing (Github Link: git@github.com:lauragails/gene_baiting.git). Bonferroni adjustments were performed and thresholds for statistical significance were delineated on the graphs. We also performed an enrichment analysis between our bait proteomes and the Grove et al., 2019 autism genome-wide association meta-analysis study ^44^ with MAGMA, as described ^107^. Mouse genes in each proximity network were mapped to human entrez gene IDs when possible, using HGNC gene nomenclature (https://www.genenames.org/, Downloaded 06/06/2023).

For cell-type specific representations, the list of genes for each bait were intersected with mouse orthologs of cell type-specific differentially expressed genes (DEGs) from a single-cell genomics study of post-mortem cortical brain tissues from autistic individuals ^8^. We used genes expressed in each cell type as the background to perform the hypergeometic test for overlap between each bait gene list and a list of autism DEGs in each cell type.

To compare HiUGE-iBioID datasets against immunoprecipitation results from brain lysate, or with recently published autism proteomic interaction datasets ^27,28^, GO analyses were conducted using input gene lists unique to each set with ShinyGO. Bait self-identifications were excluded from analyses across studies. Gene names were converted to mouse orthologs if applicable. For analyses that lack a consensus statistical domain, mouse genome was used as a non-biased background.

### Western blot

For Western blot analyses, 10μL of HiUGE-iBioID purified samples were subjected to SDS-PAGE. After transferring to nitrocellulose membranes, the blot was probed for HA-epitope (rabbit anti-HA, Cell Signaling #3724, 1:1000; or rat anti-HA, Sigma #11867423001, 1:1000), or PSD95 (mouse anti-PSD95, Abcam #ab2723, 1:1000) at 4°C overnight. Equal amounts of input protein from mouse brain lysate were also subjected to SDS-PAGE, and the blot was probed for GAPDH (rabbit anti-GAPDH, Abcam #ab9485 or Cell Signaling #2118, 1:1000) at 4°C overnight. Matching IRDye secondary antibodies (LI-COR, 1:10,000) were incubated with the blot for 1 hr at RT, and the immunosignal was detected using Odyssey FC imager (LI-COR).

### Immunocytochemistry and immunohistochemistry

Cells were fixed with 4% PFA and 4% sucrose on DIV 14, then blocked with blocking buffer (Abcam #ab126587, 1:10 in PBS with 0.3% Triton-X). Immunocytochemistry was performed by incubating with primary antibody overnight at 4°C, and with secondary antibody for 1 hr at RT. Antibodies and dilutions consisted of: mouse anti-V5-epitope (ThermoFisher #R960-25, 1:500), rabbit anti-V5-epitope (Cell Signaling #13202S, 1:1000), guinea pig anti-Homer1 (Synaptic Systems #160004, 1:2000), rabbit anti-Syngap1 (ThermoFisher #PA5-58362, 1:1000), rabbit anti-Shank2 (Cell signaling #12218S, 1:1000), mouse anti-Ank3 (ThermoFisher #33-8800, 1:1000), mouse anti-Scn2a (Antibodiesinc #75-024, 1:1000), goat anti-mouse Alexa Fluor Plus 594 (ThermoFisher #A32742, 1:1000), goat anti-guinea pig Alexa Fluor 488 (ThermoFisher #A11073, 1:1000), and goat anti-rabbit Alexa Fluor 488 (ThermoFisher #A11008, 1:1000). Cells were counterstained with 4′,6-diamidino-2-phenylindole (DAPI), mounted with FluorSave reagent (Millipore Sigma #345789), cover-slipped, and imaged on Zeiss Axio Imager M2 with Apotome module or Zeiss LSM 710 confocal microscope. For immunohistochemistry, brains were collected after intracardiac perfusion with ice cold saline and 4% PFA, cryoprotected in 30% sucrose solution, and sectioned at 40µm. Sections were blocked with blocking buffer and incubated with Alexa Fluor 594 or 568-conjugated streptavidin (ThermoFisher #S32356 or #S11226, 1:1000) overnight at 4°C. After washes, sections were counterstained with DAPI and cover-slipped for imaging on Zeiss Axio Imager M2 with Apotome module or Zeiss LSM 710 confocal microscope. Images were processed using Fiji ^108^.

### Immunoprecipitation

Expression vectors of Myc-DDK-tagged human ORF clones of ANKS1B and SYNGAP1 were purchased from Origene (#RC211877, #RC229432). V5-epitope or GFP-tagged mutants were cloned into the same expression backbone. Truncations of Syngap1 consisted of the following: N-Trunc: Δ a.a. 2-361; C1-Trunc: Δ a.a.730-1343; C2-Trunc: Δ a.a. 848-1343; C3-Trunc: Δ a.a. 1181-1343. Mutation of Syngap1 (SYNGAP1-c.2214_2217del) was introduced by site-directed mutagenesis (QuikChange Lightning, Agilent #210518). HEK293T cells were co-transfected with expression vectors using Lipofectamin 3000 (ThermoFisher). Four days following transfection, cells were lysed for Nanobody Trap experiments following the manufacturer’s protocol (Chromotek #gta-20, #gtma-20, or #v5tma-20). The bound proteins were eluted by boiling in 2X Western sample buffer and subjected to SDS-PAGE (immunoprecipitated IP fraction). The membrane was probed for Myc-epitope (Santa Cruz #sc-40, 1:250), V5-epitope (Cell Signaling #13202S, 1:1000), or GFP (Cell Signaling #2956S, 1:2000). The lysate was also subjected to SDS-PAGE (input fraction). Here, the membrane was probed for Myc-epitope (Santa Cruz #sc-40, 1:250) and GAPDH (Abcam #ab9485 or Cell Signaling #2118, 1:1000). Matching IRDye secondary antibodies (LI-COR, 1:10,000) were used to visualize immuno-signals on an Odyssey FC imager.

For immunoprecipitation from brain lysate, forebrain tissues were homogenized in T-PER protein extraction reagent (ThermoFisher #78510), cleared by centrifugation following the manufacturer’s instruction, and equally divided into replicates. The following antibodies were added to the lysate at a final concentration of 10 ug/mL: mouse anti-Anks1b (Santa Cruz #sc-376610), mouse anti-Scn2a (Antibodiesinc #75-024), or mouse control IgG (Vector labs #I-2000-1) and incubated overnight on a rotator. The next day, 25uL Pierce Protein A/G magnetic beads (ThermoFisher #88802) were added to each immunoprecipitation replicates and incubated for 1 hr under RT. Beads were washed in wash buffer containing 150mM NaCl and boiled in 2X sample buffer to elute the precipitated complexes for label-free LC-MS/MS analysis. Peptide signals were subjected to trimmed-mean normalization and summed on the protein level. Single-peptide detections and contaminants such as mouse IgG were excluded from the analysis. Enrichment was defined as Log2-FC ≥ 1 and *p*-value < 0.05 using the Duke Proteomics and Metabolomics Shared Resource (DPMSR) Proteome Discoverer Data Visualization Tool.

### AAV / Cas9-mediated expression disruption

To disrupt Syngap1 and Anks1b expression, the following genomic sequences were targeted by gRNAs using AAV: Syngap1: acggactcggtctcagcccatgg; Anks1b: attgtcccactgtttggacaggg. AAVs (PHP.eB serotype) were applied to H11-Cas9 primary neuronal cultures, with empty gRNA virus serving as negative control. Effective disruption of Syngap1 and Anks1b expression was confirmed by Western blot (Syngap1: Sigma #SAB2501893 or ThermoFisher #PA5-58362, 1:1000; Anks1b: ThermoFisher #PA5-98554, 1:1000).

### Lentiviral-mediated Scn2a CRISPRa

The all-in-one gRNA-Cas9 expression plasmid used for CRISPRa was generated by modifying the hUBC-dSpCas9-2xVP64-T2A-BSD plasmid (Addgene #162333) to remove the T2A-BSD selection marker and include a U6-gRNA scaffold. The non-targeting control gRNA and the gRNAs targeting the *Scn2a* promoter were selected from the Caprano Mouse CRISPR Activation Pooled Library ^109^ (No_current_500 (non-targeting scramble): tttttagacctaattcgcgc; Scn2a_gRNA1: cagcgattccacttgtggcc; Scn2a_gRNA2: gttgaatgttgctttgccaa; and Scn2a_gRNA3: aattacagcgattccacttg). Individual gRNAs were purchased as oligonucleotides (Integrated DNA Technologies) and cloned into the gRNA expression plasmids using BsmBI sites.

HEK293T cells were acquired from the American Tissue Collection Center (ATCC). The cells were maintained in DMEM high glucose supplemented with 10% FBS and 1x GlutaMAX Supplement (Gibco #35050061) and cultured at 37°C with 5% CO_2_. Lentivirus was produced as previously described ^110^: 16 hr before transfection, 7×10^6^ cells were plated in 12ml of transfection media (Opti-MEM I Reduced Serum Medium (Gibco #31985070), 1x GlutaMAX Supplement, 5% FBS, 1mM sodium pyruvate (Gibco #11360070), 1x MEM NEAA (Gibco #11140050) in a 10cm plate. On the day of transfection, 6mL of transfection media were removed, and the cells were transfected with Lipofectamine 3000 (Invitrogen #L3000008) and 5.4μg pMD2.G (Addgene #12259), 9.9μg psPAX2 (Addgene #12260), and 12μg of the expression vector, according to the manufacturer’s instructions. The medium was exchanged with transfection media 6hr after transfection, and the viral supernatant was harvested 24 and 48 hrs after this medium change. The viral supernatant was pooled and passed through a PVDF 0.45 μm filter (Millipore #SLHV033RB), and concentrated to 50x in 1xPBS using Lenti-X Concentrator (Clontech #631232) in accordance with the manufacturer’s protocol. The lentivirus was titered by qPCR as previously described ^111^. The concentrated lentivirus was snap-frozen and stored at −80°C as single-use aliquots.

To assess the effect of CRISPRa transcriptional activation, cultured neurons were treated with Scn2a-CRISPRa or non-targeting control lentiviral vectors at DIV0. On DIV11, the cDNA was prepared using Cells-to-cDNA kit (ThermoFisher #AM1723) following the manufacturer’s instructions. Predesigned KiCqStart SYBR green primers (Sigma #KSPQ12012) targeting mouse Scn2a and Actb were used for qPCR experiments with PowerUp SYBR green master mix (ThermoFisher #A25742). Specific on-target amplifications were confirmed by gel electrophoresis and TOPO sequencing of the PCR products (ThermoFisher #450030). The gRNA2 was selected for the experiments in this study based upon its superior ability to upregulate Scn2a over gRNA1 and gRNA3 in cultured neurons.

### AAV-mediated expression of SCN1B and FGF12

To express additional SCN1B or FGF12, cDNAs (Origene #RC209565, #RC215868) were cloned into an AAV-expression vector downstream of the hSyn promotor. A non-targeting AAV-Flex-GFP vector (from Dr. Il Hwan Kim) was used as a negative control. Cultured neurons were treated with these AAV vectors (PHP.eB serotype) at DIV0 and lysed for Western blot analysis on DIV11. Effective expression was confirmed by Western blot (SCN1B: Cell Signaling #13950S, 1:1000; FGF12: Proteintech #13784-1-AP, 1:1000).

### Microelectrode Array (MEA)

48-well MEA plates (Axion Biosystems #M768-KAP-48 or #M768-tMEA-48W) were coated with 1 mg/mL poly-L-lysine in borate buffer (pH 8.5). Forebrain tissue was rapidly isolated from P0-1 mice, dissociated with papain, and spotted at a density of 150,000 cells per the inner growth area of each well. For experiments that required genotyping, brain tissue was temporarily stored in Hibernate A solution (ThermoFisher #A1247501) at 4°C following the manufacturer’s instructions until ready for dissociation and plating. Viral treatments were applied at the day of plating. Recordings were conducted on a Maestro MEA system (Axion Biosystems) on DIV 8, 11, and 14 for 10 min for each session, after 10 min of acclimation to the recording chamber (37°C, 5% CO_2_ environment). After each recording, a half-change of growth media (Neurobasal A supplemented with B27, GlutaMax, and 10 µg/mL Gentamicin) was performed. Recording data were analyzed using the Axion Neural Metric Tool. Single electrode bursts were defined as a minimum of 5 spikes, separated by an inter-spike interval (ISI) of no more than 100 msec. Network bursts were defined as a minimum of 10 spikes, separated by an ISI of no more than 100 msec with at least 25% of the electrodes active. Statistical analyses (one-way ANOVA followed by pair-wise Tukey HSD *post-hoc* tests) were performed using JMP Pro (SAS).

### Behavioral Tests

#### Elevated zero maze

Anxiety-like behaviors were assessed in the elevated zero maze as described under 40-60 lux illumination ^112^. Mice were housed overnight in the test room and tested individually the next day. Animals were placed into a closed area of the maze and provided 5 min of free exploration. Videos were scored by trained observers blinded to the genotype and sex of the animals using the Noldus Observer XT15 program (Leesburg, VA) for the percent time in the open areas and the distance traveled.

#### Hole-board test for repetitive behaviors

Mice were housed in the test room overnight and the next day were examined in the hole-board test as described ^112^. Individual mice were placed into a 42 x 42 x 30 cm open field (Omnitech Electronics, Columbus, OH) and given free exploration of the apparatus for 10 min under 180 lux illumination. The hole-board apparatus consisted of a white Plexiglas floor with 16 equally spaced holes (3 cm in diameter) arranged in 4 rows. Head-dips into the holes were filmed and the videos were scored with the TopScan program (CleverSys, Reston, VA) for the numbers of head-dips and the location of each head-dip.

#### Self-grooming

Individual mice were housed in the test room overnight and the next day were habituated to clean home-cages for 10 min prior to testing ^113^. Mice were filmed for 10 min for self-induced grooming. Subsequently, the videos were converted for analysis using Noldus MediaRecorder2. Grooming behavior was scored using TopScan software (CleverSys, Reston, VA) for the duration of self-grooming.

#### Ultrasonic vocalizations (USVs)

Male WT and Scn2a^+/R102Q^ mice were housed individually 1 week before testing. Initially, males were primed twice with soiled bedding from estrous C57BL/6J females for 2 days. Subsequently, males were placed on clean bedding for 1 day, then on soiled bedding for another 2 days, and finally were returned to clean bedding overnight. Males were tested the next day. They were acclimated to the recording chambers for 2 min and then were introduced to an unfamiliar C57BL/6J female (12-16 weeks of age) for 8 min. Ultrasonic calls were recorded over the entire 10 min test as waveform audio files. The data were analyzed by scorers blinded to the genotypes of the mice using Avisoft SASLab Pro (Glienicke/Nordbahn, Germany) for the numbers of calls, call duration, and USV frequencies.

#### Social behavior tests

Social behavior in male WT and Scn2a^+/R102Q^ mice was assessed in the resident-intruder and social dyadic assays ^114^. The Scn2a males were housed individually on the same bedding for 14 days prior to testing; the partner C3H/HeJ mice (Jackson Laboratory, Bar Harbor, ME) were group-housed at 3-4 mice/cage. Both social tests were conducted with C3H/HeJ males matched for body weight with the Scn2a males. Note, the C3H/HeJ mice provide high amounts of social investigation without initiating unprovoked agonistic or attack behaviors ^115^. Social testing began within two hr after the lights had extinguished and was conducted under red light. Prior to testing, the chow, water bottle, and metal frame holding the water bottle were removed. The filter was removed from the top of the cage so behaviors could be filmed and the mouse could not escape. Tests were terminated if attacks by either Scn2a or C3H/HeJ males occurred for more than 1 min or if a mouse was injured. All behaviors were tested in the Social StereoScan apparatus (Cleversys) where behaviors can be filmed simultaneously from the top and sides of the test chamber. All videos were scored using Noldus Observer (version 15) by trained personnel blinded to the genotypes of the mice.

Mice were tested first in the resident-intruder assay. Here, a C3H/HeJ male was introduced into the home-cage of the WT or Scn2a^+/R102Q^ male. Animals were allowed to interact for 5 min, after which the C3H/HeJ male was removed to its home-cage. Following this testing, the cages housing the individual Scn2a mice were replaced with new cages and bedding. Seven days later dyadic testing began where a WT or Scn2a^+/R102Q^ male was paired with an unfamiliar C3H/HeJ male in a novel clear Plexiglas chamber (32 x 20 x 20 cm). Mice were placed at opposite ends of the chamber divided in half with a white polyfoam partition. Following a 5 min habituation, the partition was removed and the mice were permitted to interact for 5 min. All behaviors in both social tests were calculated as a rate per 5 min [(numbers of incidences/total test time in sec) x 300 sec] and were analyzed over four response domains ^114^. Mild-social investigation consisted of approaching, sniffing or nosing (ano-genital region), climbing onto the side or back, and/or grooming the face of the partner. Reactivity referred to boxing, holding, kicking, startling, and/or jumping in the presence of the partner. Agonistic behaviors denoted tail rattling to the partner and/or feinting, chasing, lunging, biting, climb-grooming, and/or attacking the partner. Withdrawal behaviors (no acknowledgement of partner) included walking-away, turning-away (without leaving proximity of the partner), or digging in the bedding when the partner contacts or attempts to interact.

#### Statistics

The data were presented as means and standard errors of the mean (SEMs). The zero maze, hole-board, self-grooming, and USV data were analyzed by independent-samples t-tests and the social behavior data were analyzed by repeated-measures ANOVA followed by Bonferroni corrected pair-wise comparisons. A *p* < 0.05 was considered significant. All statistical analyses were performed with IBM SPSS Statistics 28 programs (IBM, Chicago, IL) and the data were graphed using GraphPad Prism (Boston, MA).

## Supplementary Materials

**Fig. S1.**
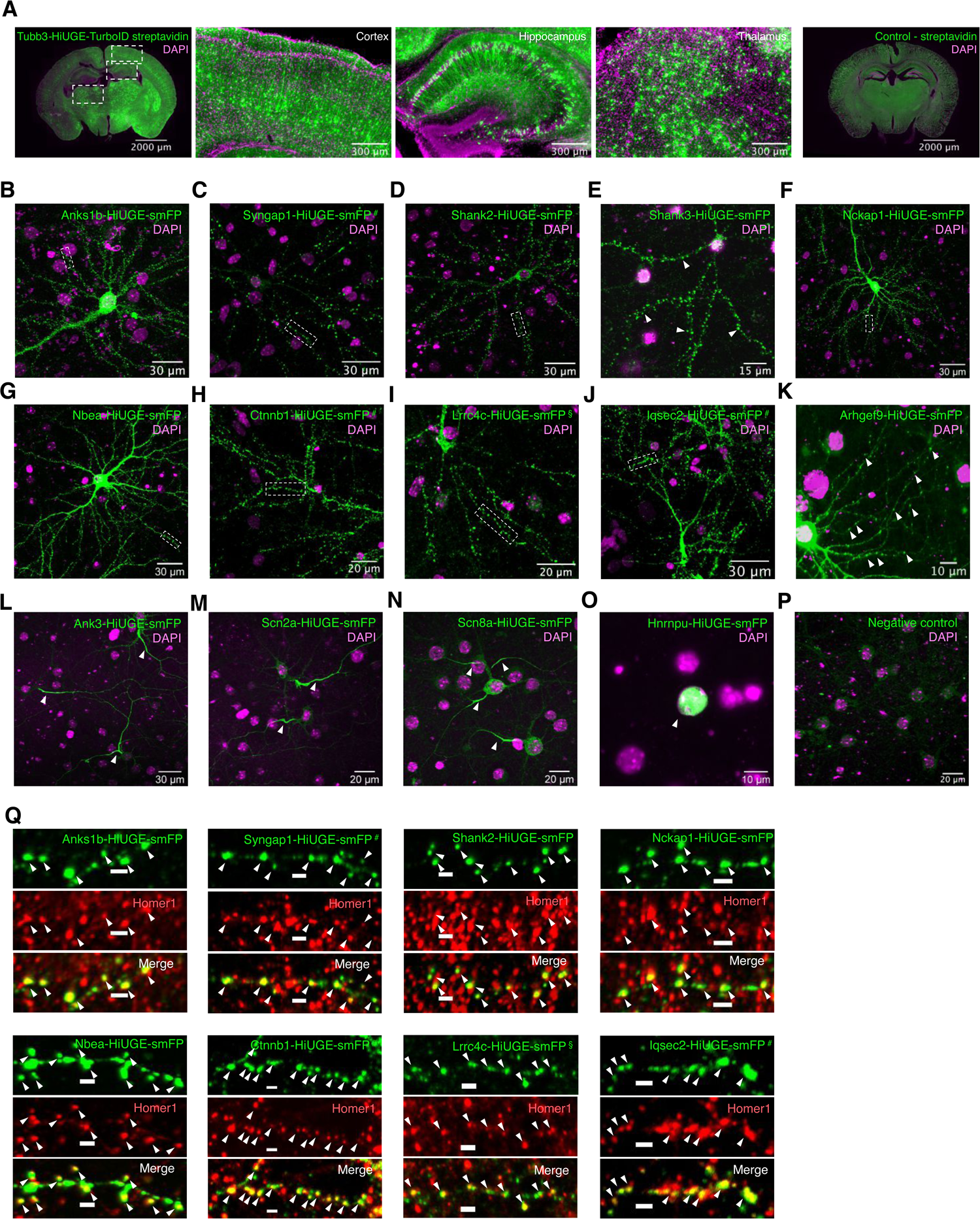
HiUGE labeling of 14 high-risk autism proteins. (**A**) Representative images showing wide-spread TurboID-mediated biotinylation across the brain following labeling Tubb3 with HiUGE. (**B-O**) Representative images showing correct localization (arrowheads) of 14 high-risk autism proteins labeled with a highly antigenic “spaghetti monster” fluorescent protein (smFP), with boxed regions enlarged in (**Q**) to show colocalization with a synaptic marker Homer1. PDZ-binding motifs were preserved using #: intron-targeting strategy and §: modified donor incorporating native PDZ-binding motif. (**P**) Representative image of the negative control. Scale bar in the enlarged view represents 2μm.

**Fig. S2.**
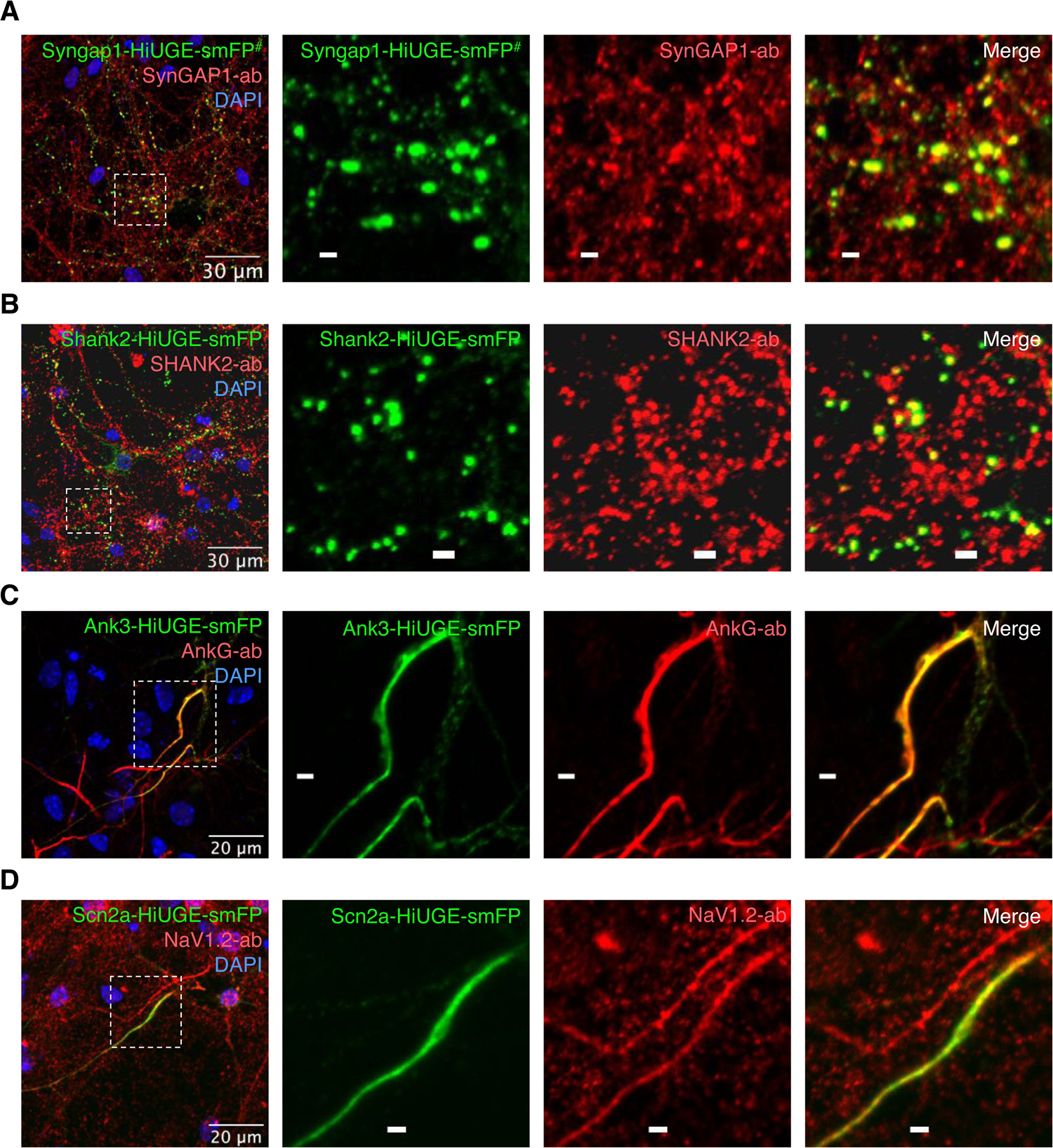
Immunofluorescence detection of representative bait proteins with HiUGE labeling. (**A-D**) Representative images showing colocalization of smFP-labeled proteins with immunofluorescence detected by specific antibodies against native proteins (Syngap1, Shank2, Ank3, Scn2a). Scale bar in the enlarged view represents 2μm.

**Fig. S3.**
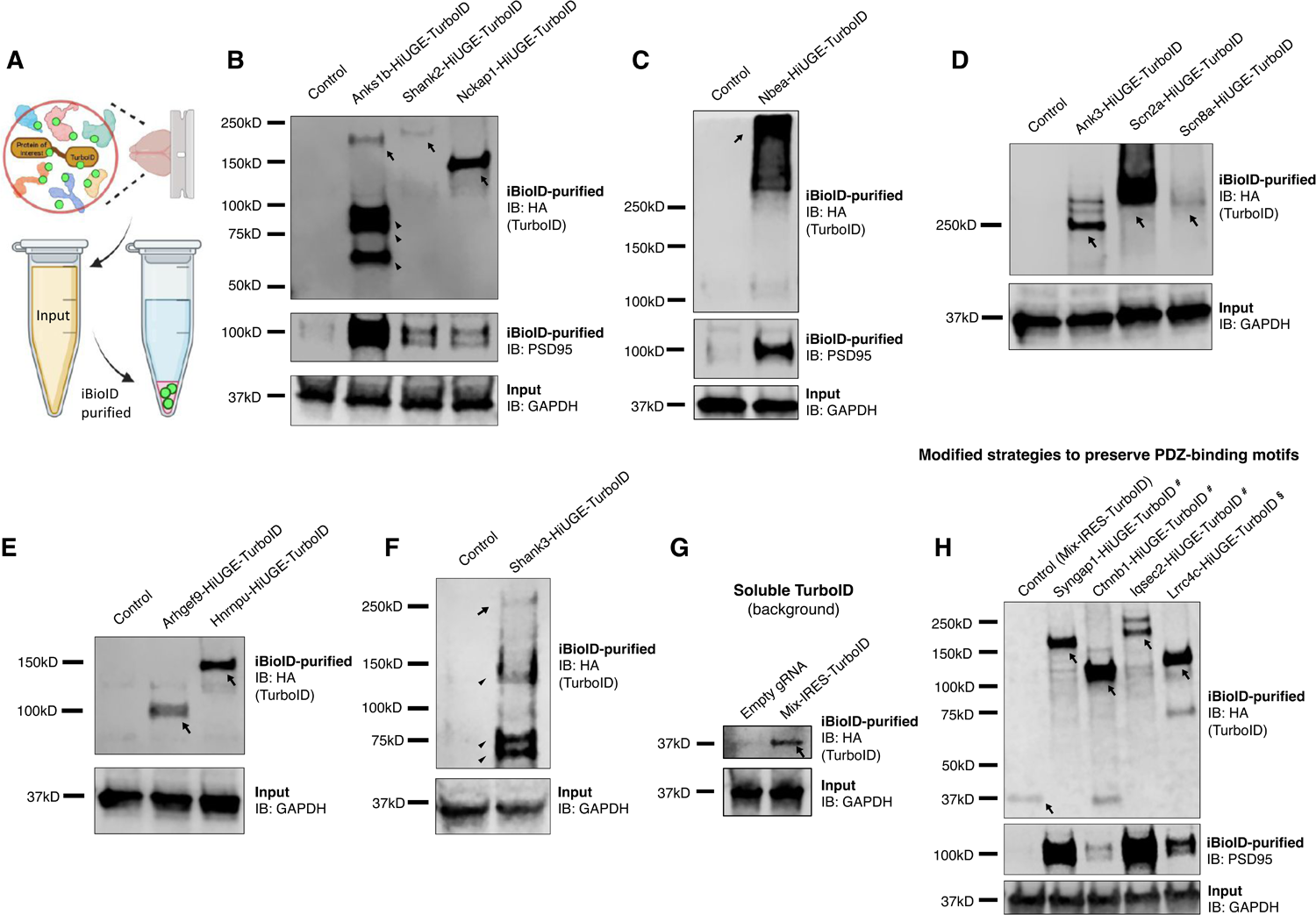
Western blot of HiUGE-iBioID purified samples following streptavidin pulldown. (**A**) Schematic illustration of enriching biotinylated proteins by streptavidin pulldown. (**B-H**) Western blot images showing detection of TurboID-HA fusion proteins at the expected molecular masses following purifications, (**B, C, H**) detection of an expected synaptic interactor (PSD95) is also confirmed. (**G, H**) Western blot images showing detection of soluble TurboID-HA (as a survey for background) by multiplexed insertion of IRES-TurboID-HA donor near 3’UTR.

**Fig. S4.**
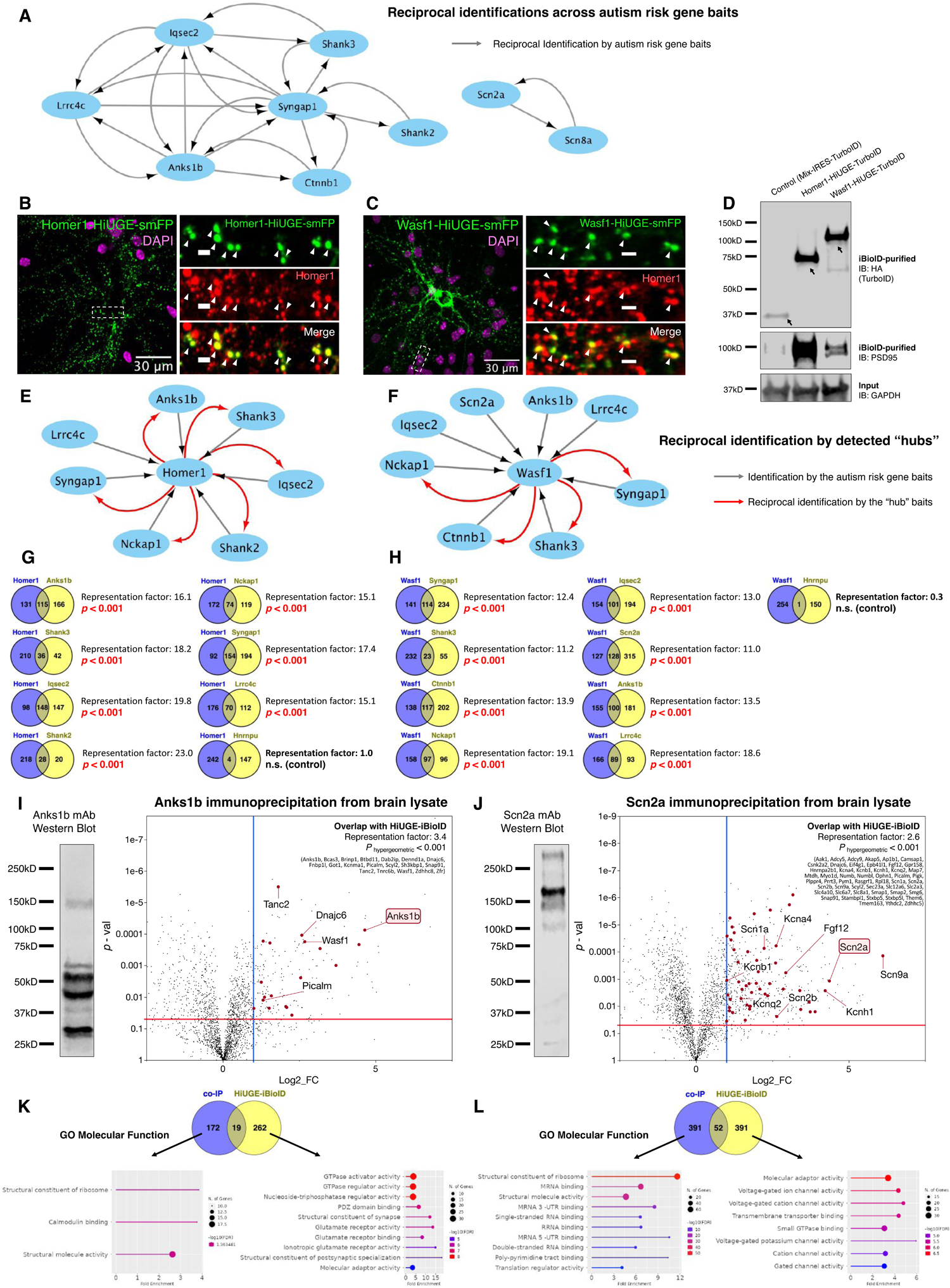
Reciprocal validation and comparison with immunoprecipitation. (**A**) Network graph showing extensive reciprocal identifications amongst autism risk protein baits. (**B-D**) Immunofluorescence and Western blot validation of HiUGE-iBioID for Homer1 and Wasf1, two exemplary proteomic “hubs” detected by many baits. (**E, F**) Network graphs showing reciprocal identifications of the initial baits by these “hubs”. (**G, H**) Hypergeometric analyses of overlap between the HiUGE-iBioID proteomes of the initial baits and the “hubs” show highly significant overlaps. No significant overlap is detected when comparing to the Hnrnpu dataset. **(I, J)** Comparison of HiUGE-iBioID results with proteomic detections following immunoprecipitation using monoclonal antibodies. Western blot images of brain lysate samples using these antibodies are shown. Genes that overlap with HiUGE-iBioID are highlighted on the volcano plot, and a few genes related to synaptic and ion channel functions are labeled. Significance of overlap is determined by hypergeometric test. (**K, L**) Comparative GO analyses of the gene sets that are exclusive to either the immunoprecipitation or the HiUGE-iBioID datasets. The statistical domain is the cumulative proteomic detections of brain-derived samples in our lab (Table S5).

**Fig. S5.**
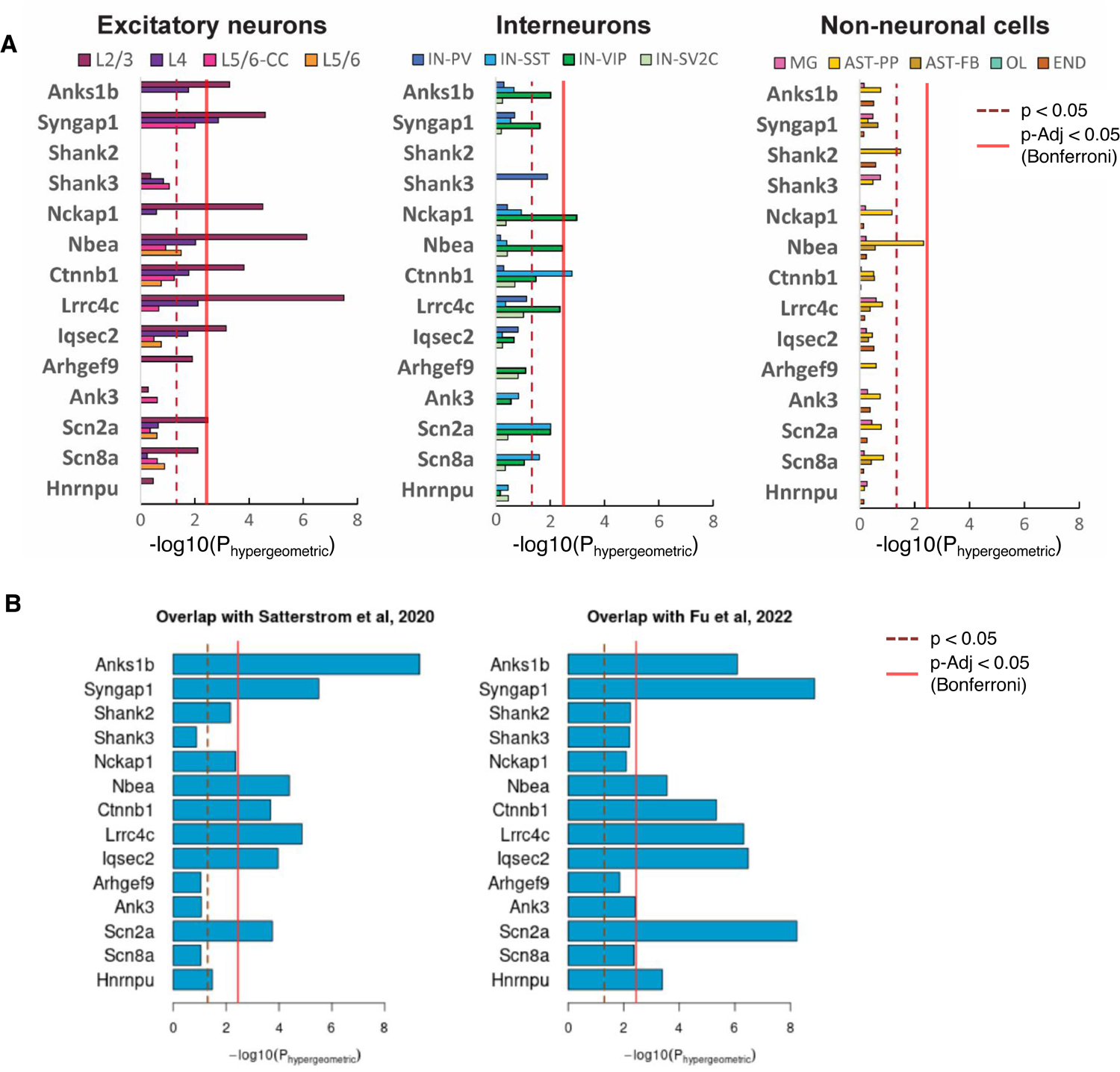
Overlap of HiUGE-iBioID interactomes with published datasets. (**A**) Significance of interactome overlap with differentially expressed genes (DEGs) found in autistic individuals across excitatory neurons, interneurons, and non-neuronal cell populations. (**B**) Significance of interactome overlap with autism risk genes identified by Satterstrom et al. ^3^, and Fu et al. ^7^. Thresholds for statistical significance were delineated on the graphs.

**Fig. S6.**
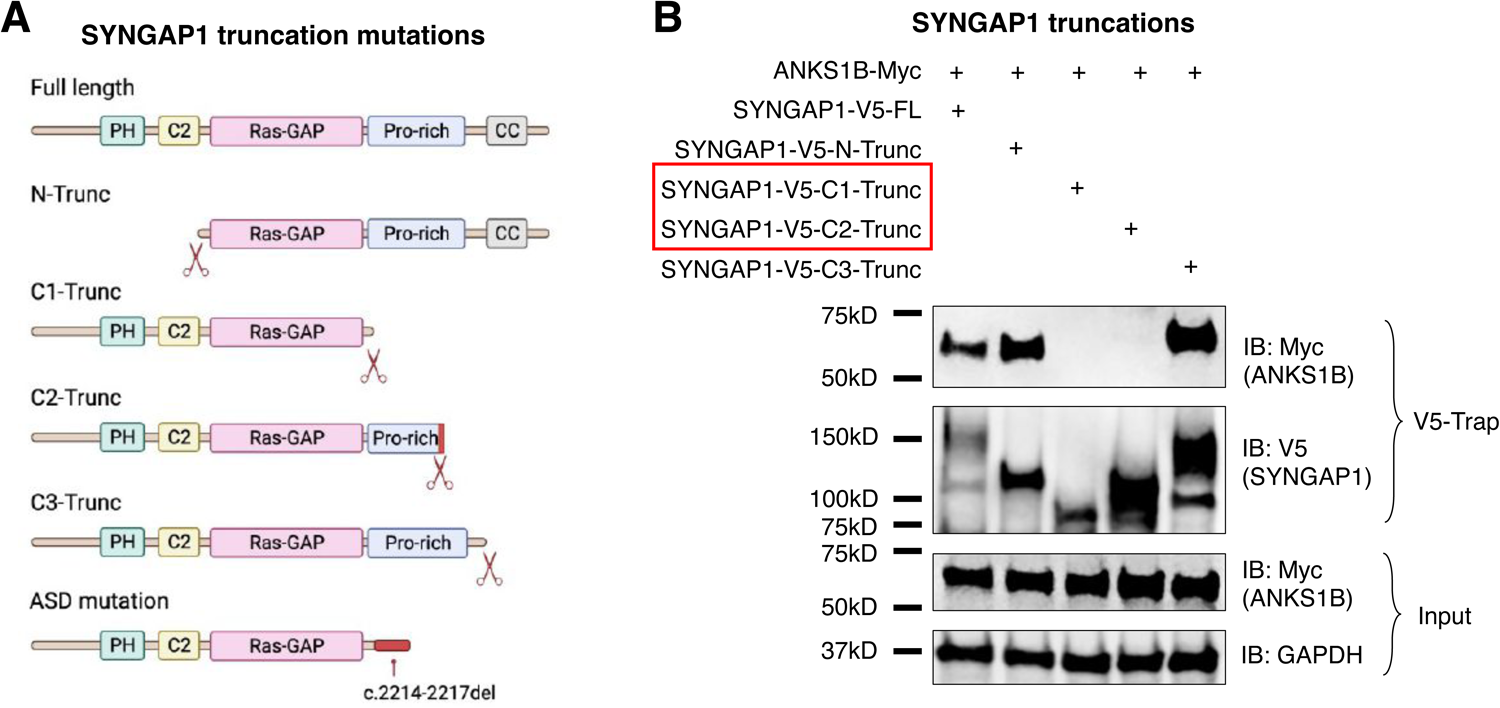
Structure-function analysis of SYNGAP1-ANKS1B interaction. (**A**) Schematic illustration of assessing the ANKS1B interaction with SYNGAP1 truncations using human cDNA constructs expressed in HEK293T cells. (**B**) Co-immunoprecipitation results showing the interaction with ANKS1B ablated in C1- and C2- SYNGAP1 truncations while retained in C3- and N- truncations.

**Fig. S7.**
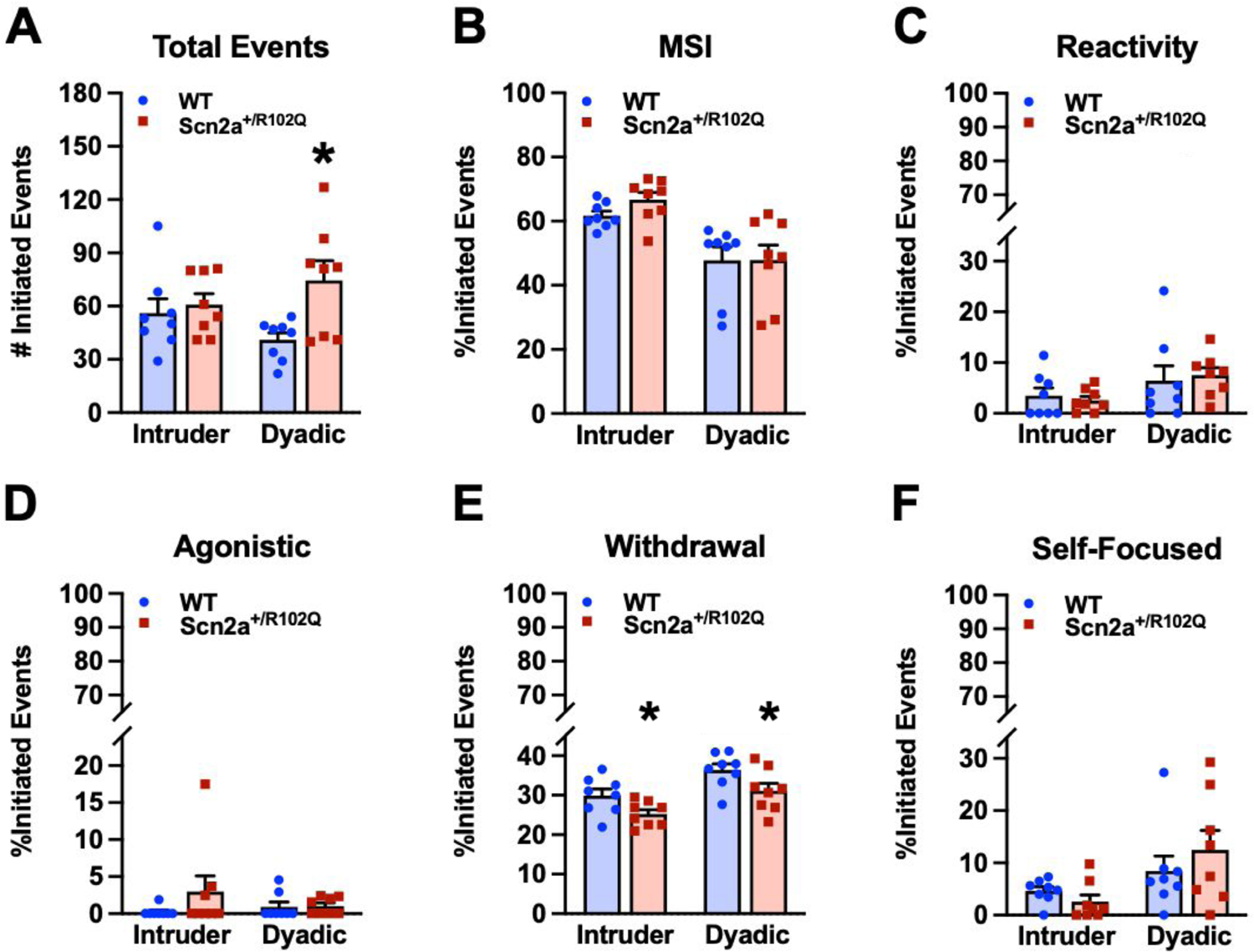
Additional social behavior data for the Scn2a^+/R102Q^ mice. (**A**) Scn2a^+/R102Q^ males initiated more overall social events in the dyadic assay with C3H/HeJ males than WT males (*p* = 0.013). (**B**) No genotype effect was detected for the percent of mild social interactions (MSI). (**C**) No genotype effect was detected for the percent of reactivity events. (**D**) The numbers of agonistic events were very low and were not distinguished by social test or genotype. (**E**) The percent of withdrawal events from C3H/HeJ partners was lower in the Scn2a^+/R102Q^ males than WT males (genotype effect, *p* = 0.020). (**F**) No genotype effect was detected for the self-focused behaviors. The data are presented as means ± SEMs and were analyzed with RMANOVA with Bonferroni corrections, n = 8 mice / genotype. *: *p* < 0.05, WT vs. Scn2a^+/R102Q^ mice. Additional statistics are summarized in Table S6.

**Fig. S8.**
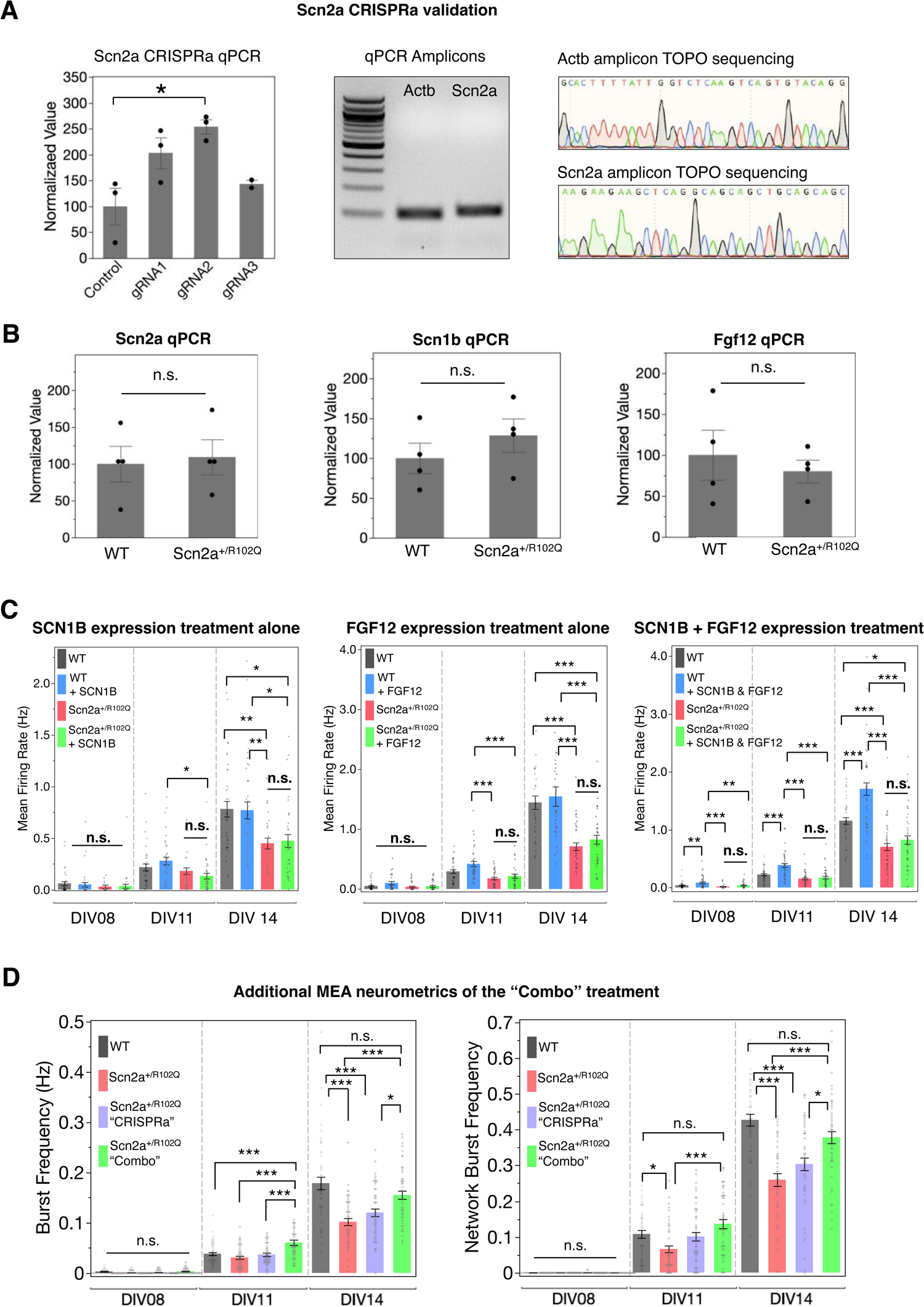
>Additional data of the Scn2a^+/R102Q^ phenotypic rescue experiment. (**A**) Quantitative PCR screening of three different CRISPRa gRNAs targeting Scn2a *in vitro*. The gRNA2 shows the best performance in upregulating Scn2a expression, and is used for subsequent experiments (ANOVA followed by Dunnett’s test, n = 3 wells). Specificity of the qPCR assay is validated by sequencing the amplicon. (**B**) mRNA expression levels of Scn2a, Scn1b, and Fgf12 in cultured Scn2a^+/R102Q^ mutant neurons are comparable to that of WT (two-tailed *t*-test, n = 4 wells). (**C**) MEA neurometrics following overexpression of SCN1B, FGF12, or combined showing ineffective rescue of the Scn2a^+/R102Q^ phenotype, One-way ANOVA followed by *post-hoc* Tukey HSD tests (n= 36 or 48 wells). (**D**) Additional neurometrics from the MEA recording of the “Combo” treatment experiment. One-way ANOVA followed by *post-hoc* Tukey HSD tests (n= 48 wells). *: *p* < 0.05; **: *p* < 0.01; ***: *p* < 0.001; n.s.: non-significant. Plots are mean ± SEM.

**Fig. S9.**
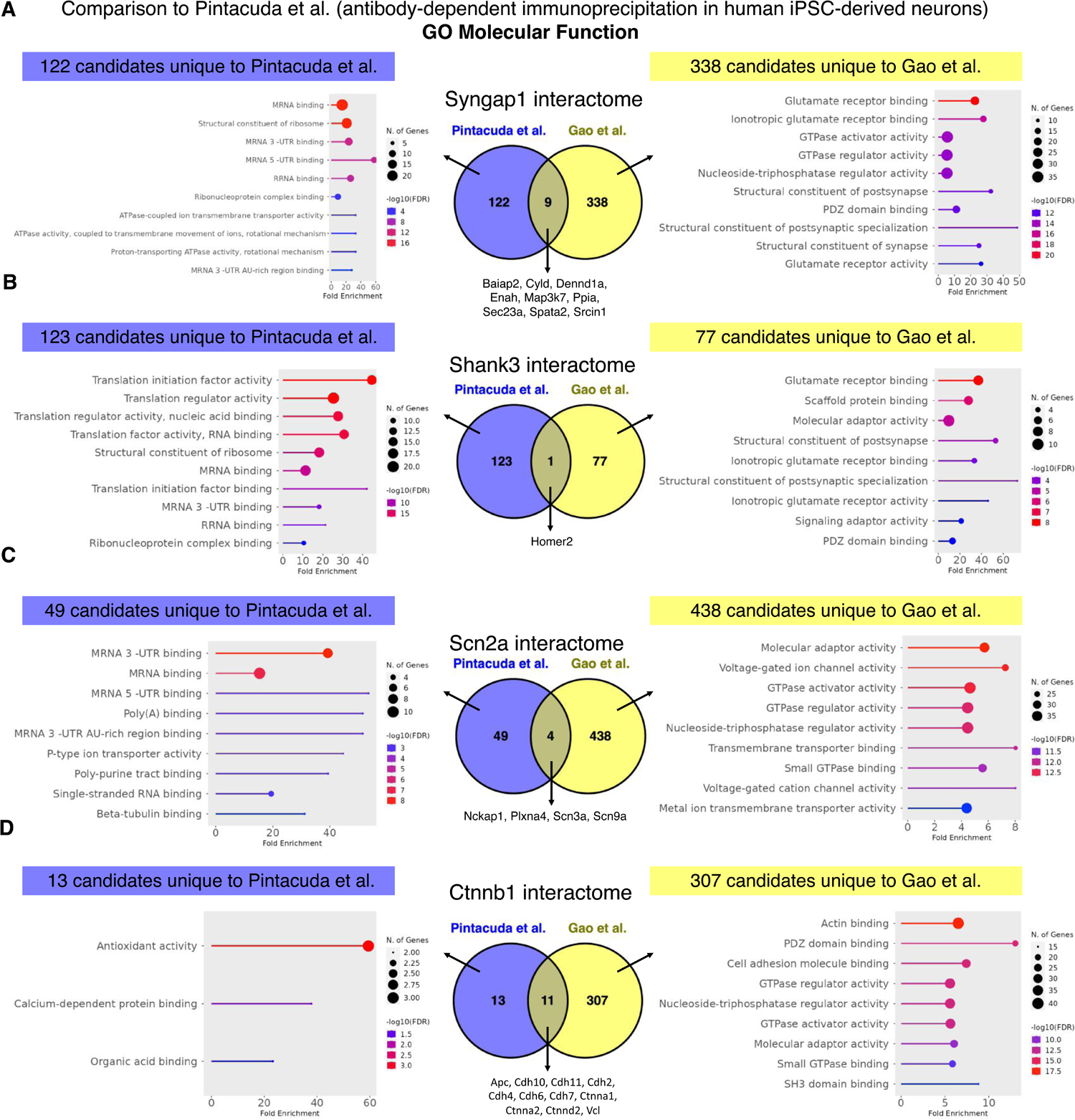
Comparative GO analysis with a recently published autism risk protein immunoprecipitation dataset using human iPSC-derived neurons. (**A-D**) Comparative GO analyses of genes that are exclusive to either the immunoprecipitation dataset using human iPSC-derived neurons ^28^ or the HiUGE-iBioID dataset for Syngap1, Shank3, Scn2a and Ctnnb1. The human genes were converted to mouse orthologs before intersecting with HiUGE-iBioID dataset. Bait self-IDs were excluded for this analysis. Due to the lack of a consensus statistical domain, mouse genome was used as a non-biased background. Top molecular function GO terms are shown.

**Fig. S10.**
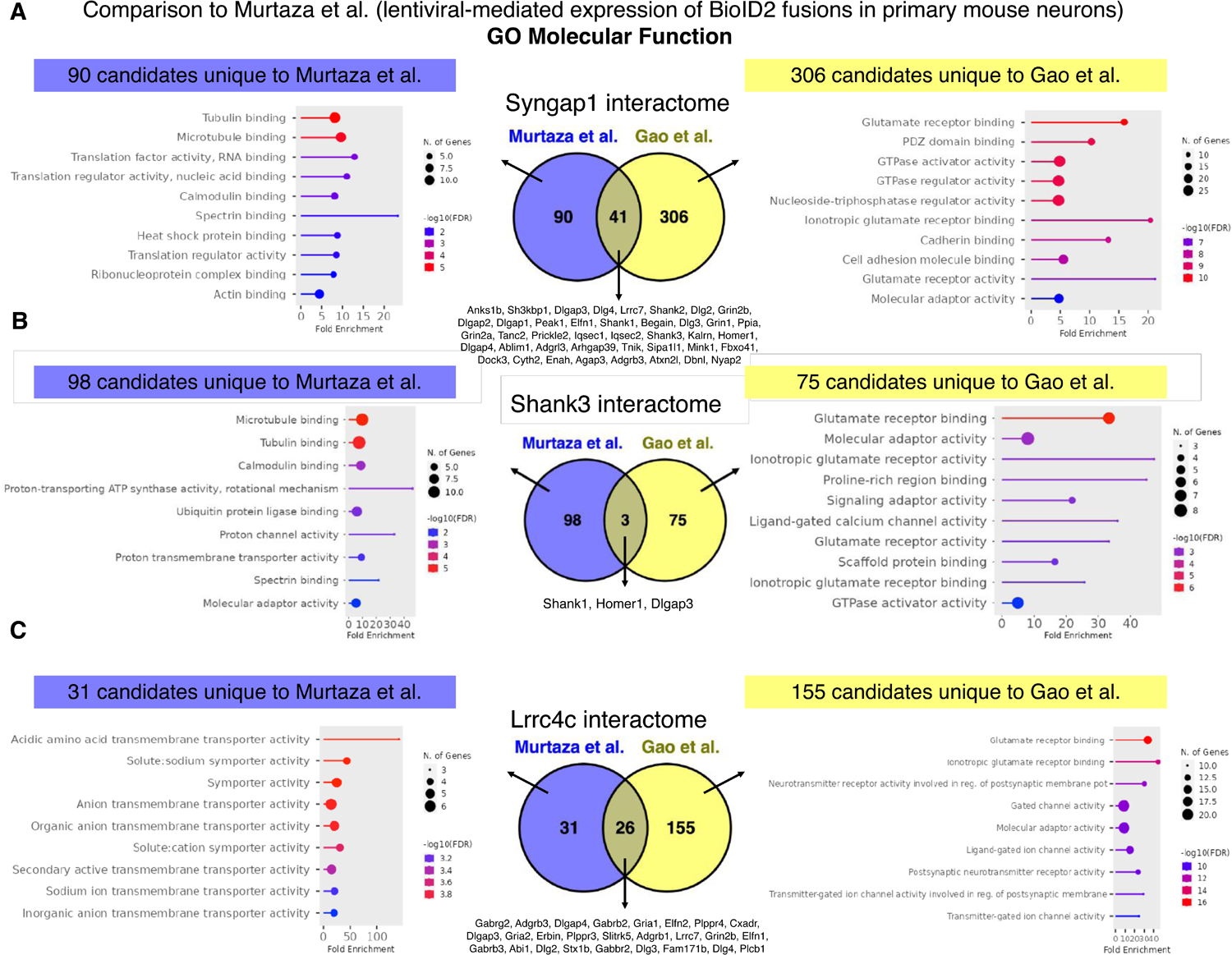
Comparative GO analysis with a recently published autism risk protein recombinant BioID dataset using cultured neurons. (**A-C**) Comparative GO analyses of genes that are exclusive to either the recombinant BioID expression dataset using primary mouse neurons ^27^ or the HiUGE-iBioID dataset for Syngap1, Shank3, and Lrrc4c. Bait self-IDs were excluded for this analysis. Due to the lack of a consensus statistical domain, mouse genome was used as a non-biased background. Top molecular function GO terms are shown.

**Fig. S11.**
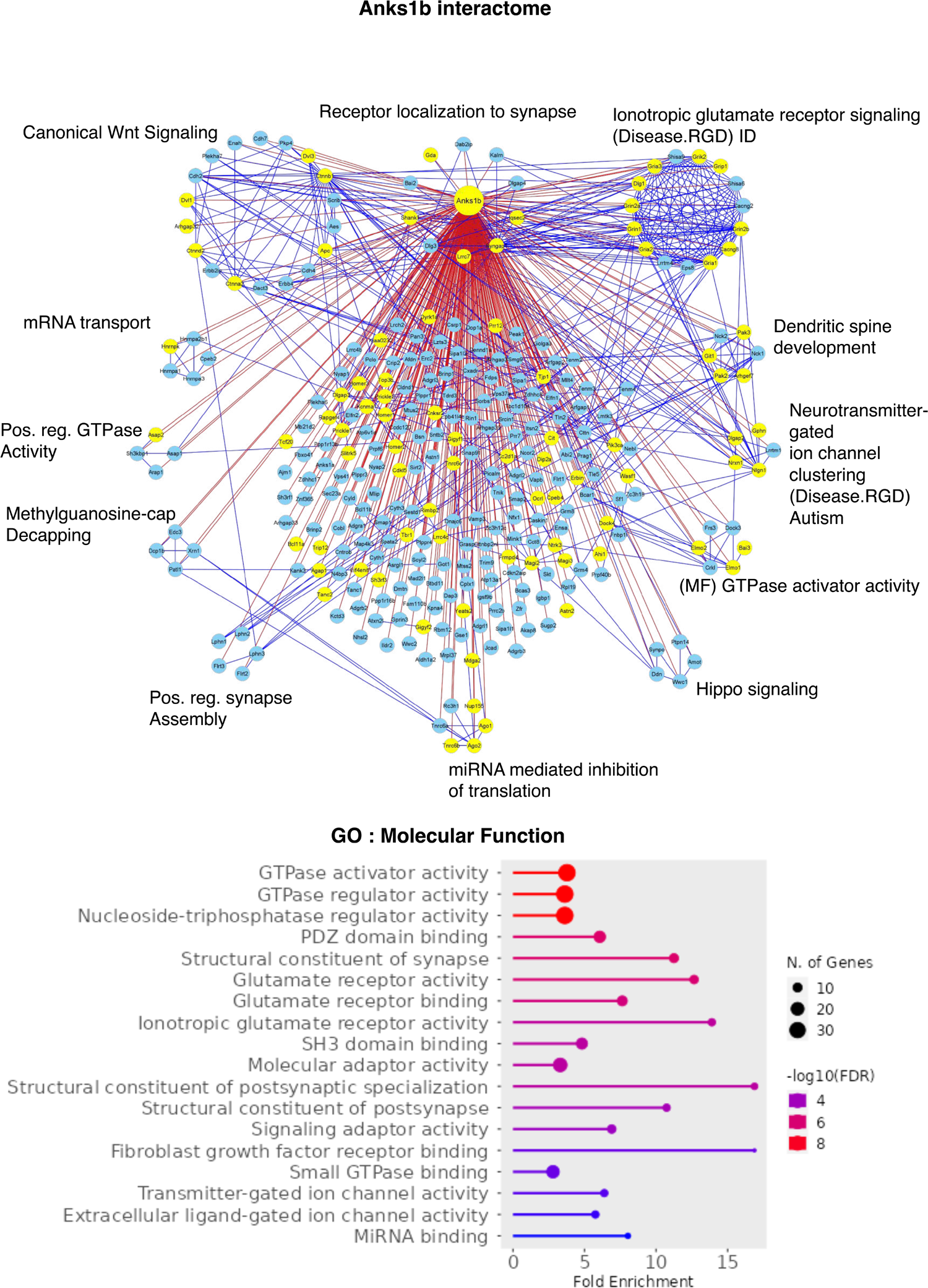
Individual interactome networks of 14 autism risk proteins and the additional 2 “hub” proteins. Interactome network associated with the bait protein is shown in each Figure. Blue lines denote STRING interactions and red lines signify identified HiUGE-iBioID interactions. Yellow nodes highlight proteins encoded by SFARI gene orthologs. Annotations denote exemplary significant gene ontology (GO) terms associated with the protein clusters segregated by MCL. Unless otherwise specified, the Biological Process pathway database was used. CC: Cellular Component pathway database. Charts of GO results of the overall bait interactomes using the Molecular Function (MF) pathway database are shown as well, below each network plot.

**Fig. S12.**
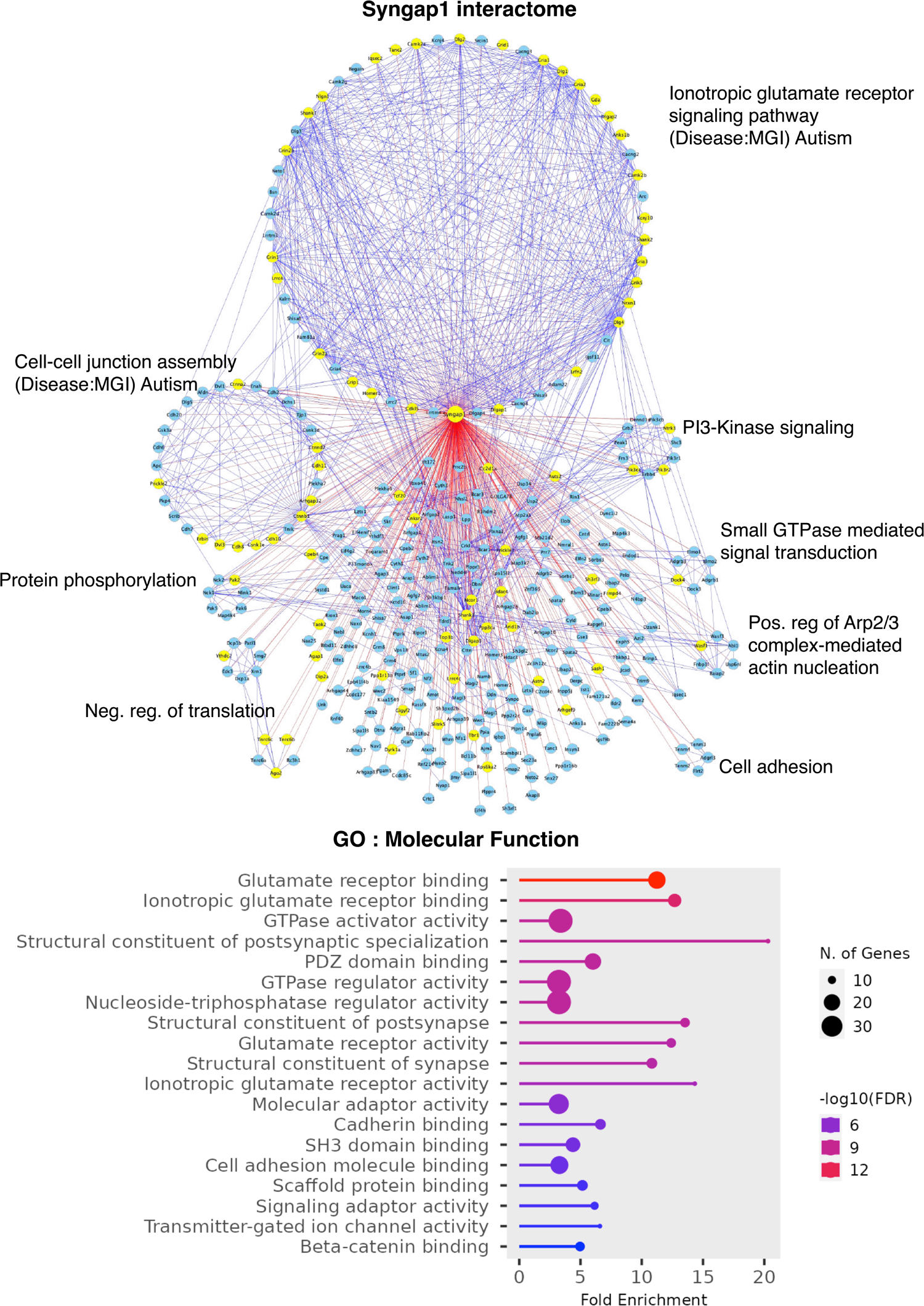
Individual interactome networks of 14 autism risk proteins and the additional 2 “hub” proteins. Interactome network associated with the bait protein is shown in each Figure. Blue lines denote STRING interactions and red lines signify identified HiUGE-iBioID interactions. Yellow nodes highlight proteins encoded by SFARI gene orthologs. Annotations denote exemplary significant gene ontology (GO) terms associated with the protein clusters segregated by MCL. Unless otherwise specified, the Biological Process pathway database was used. CC: Cellular Component pathway database. Charts of GO results of the overall bait interactomes using the Molecular Function (MF) pathway database are shown as well, below each network plot.

**Fig. S13.**
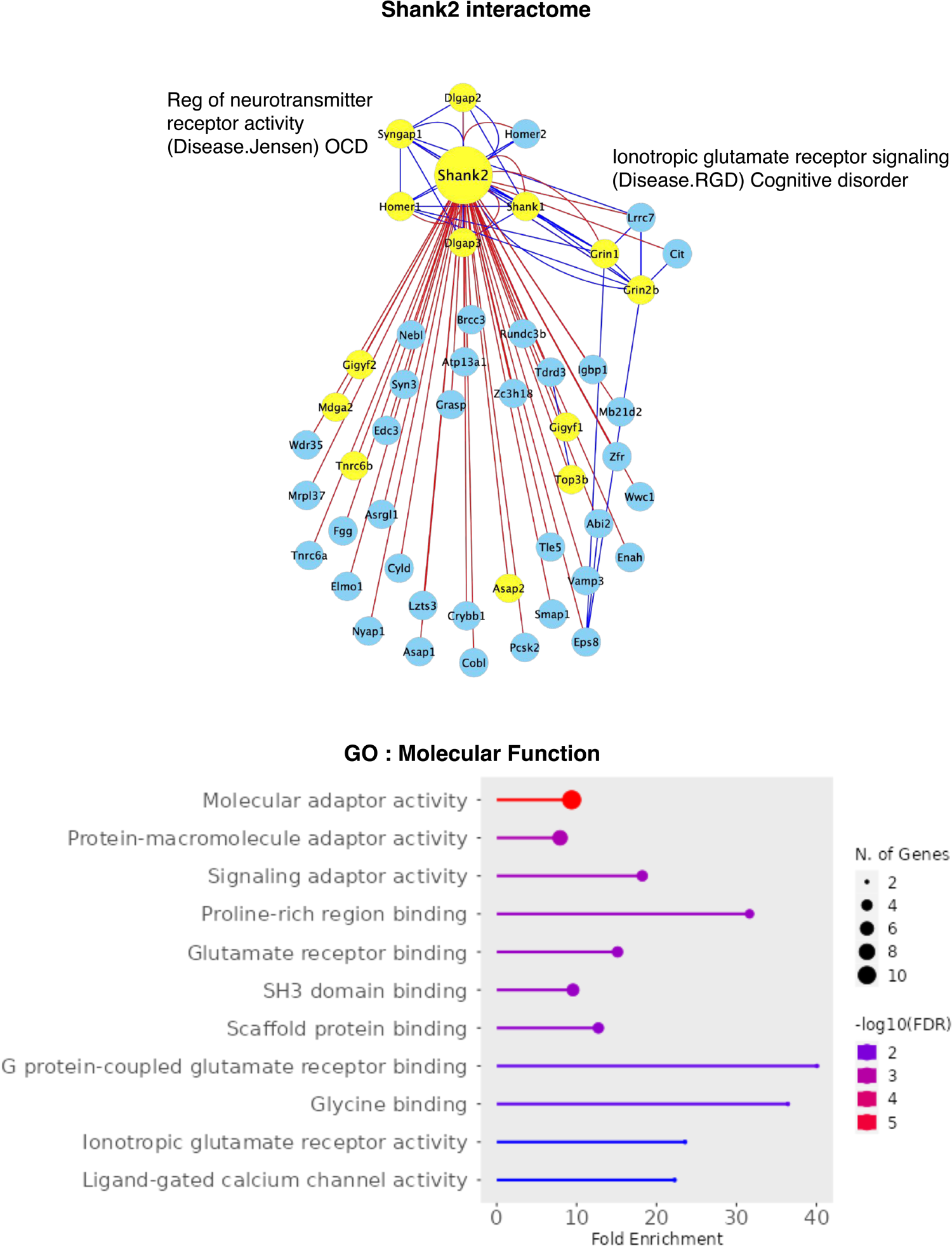
Individual interactome networks of 14 autism risk proteins and the additional 2 “hub” proteins. Interactome network associated with the bait protein is shown in each Figure. Blue lines denote STRING interactions and red lines signify identified HiUGE-iBioID interactions. Yellow nodes highlight proteins encoded by SFARI gene orthologs. Annotations denote exemplary significant gene ontology (GO) terms associated with the protein clusters segregated by MCL. Unless otherwise specified, the Biological Process pathway database was used. CC: Cellular Component pathway database. Charts of GO results of the overall bait interactomes using the Molecular Function (MF) pathway database are shown as well, below each network plot.

**Fig. S14.**
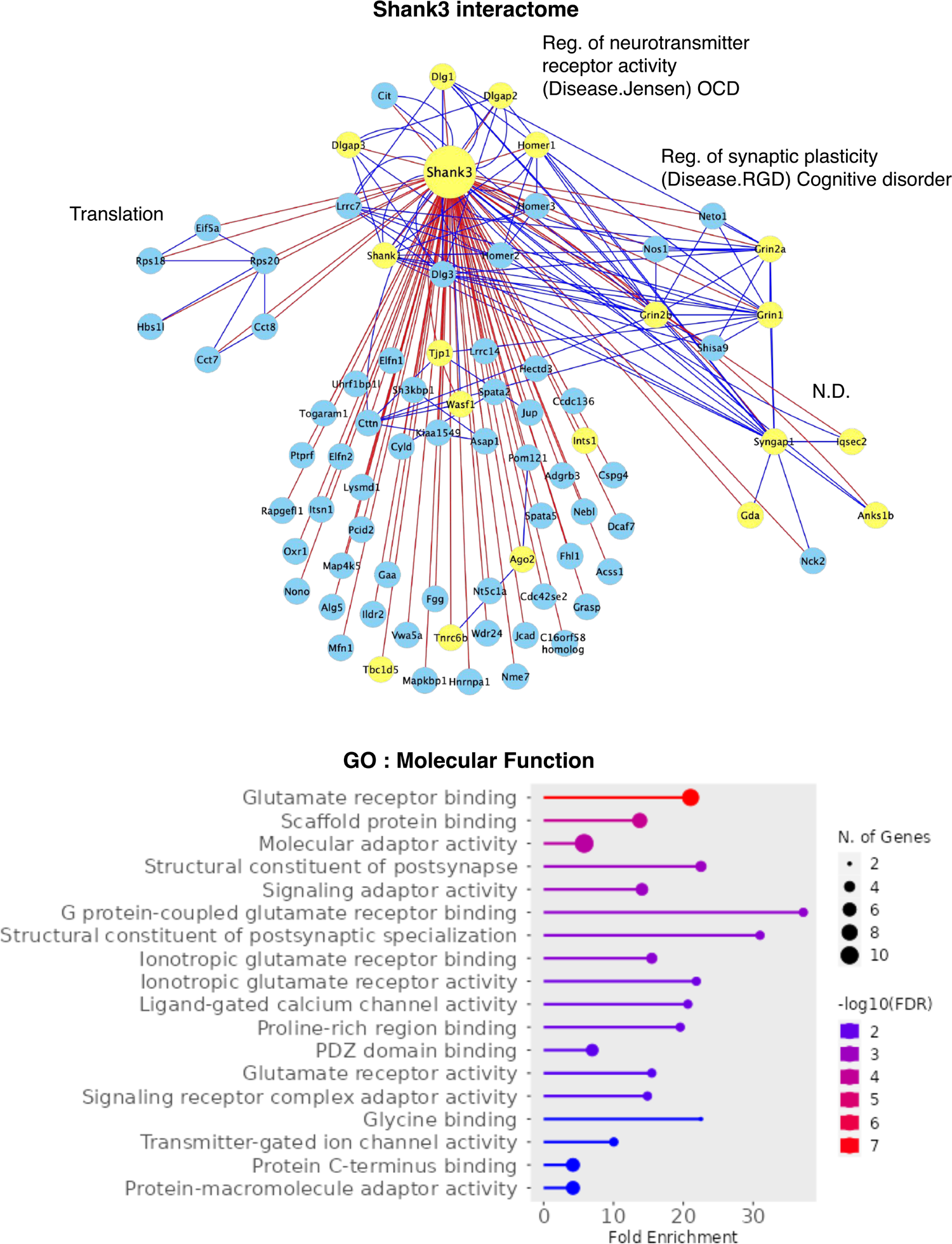
Individual interactome networks of 14 autism risk proteins and the additional 2 “hub” proteins. Interactome network associated with the bait protein is shown in each Figure. Blue lines denote STRING interactions and red lines signify identified HiUGE-iBioID interactions. Yellow nodes highlight proteins encoded by SFARI gene orthologs. Annotations denote exemplary significant gene ontology (GO) terms associated with the protein clusters segregated by MCL. Unless otherwise specified, the Biological Process pathway database was used. CC: Cellular Component pathway database. Charts of GO results of the overall bait interactomes using the Molecular Function (MF) pathway database are shown as well, below each network plot.

**Fig. S15.**
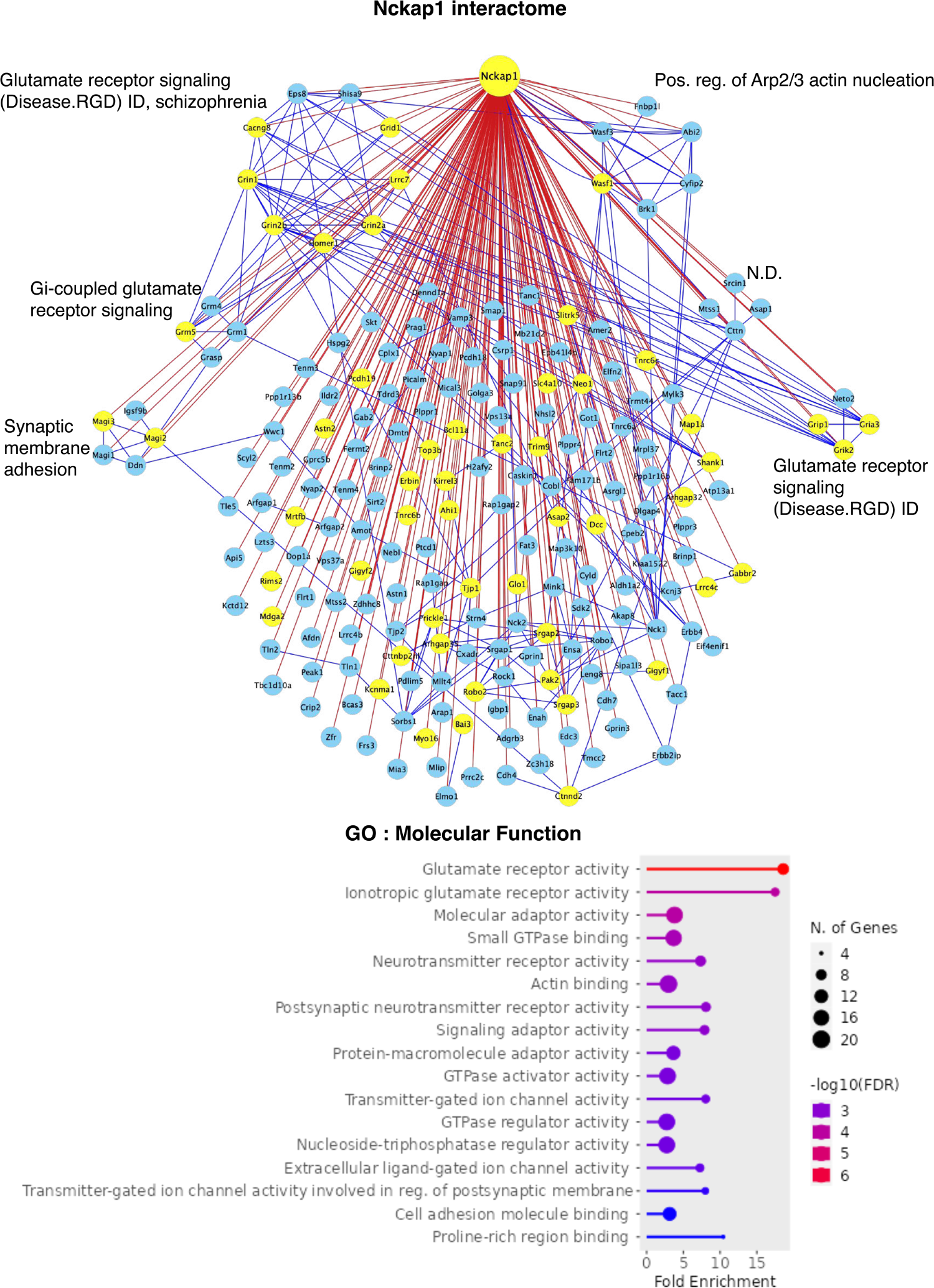
Individual interactome networks of 14 autism risk proteins and the additional 2 “hub” proteins. Interactome network associated with the bait protein is shown in each Figure. Blue lines denote STRING interactions and red lines signify identified HiUGE-iBioID interactions. Yellow nodes highlight proteins encoded by SFARI gene orthologs. Annotations denote exemplary significant gene ontology (GO) terms associated with the protein clusters segregated by MCL. Unless otherwise specified, the Biological Process pathway database was used. CC: Cellular Component pathway database. Charts of GO results of the overall bait interactomes using the Molecular Function (MF) pathway database are shown as well, below each network plot.

**Fig. S16.**
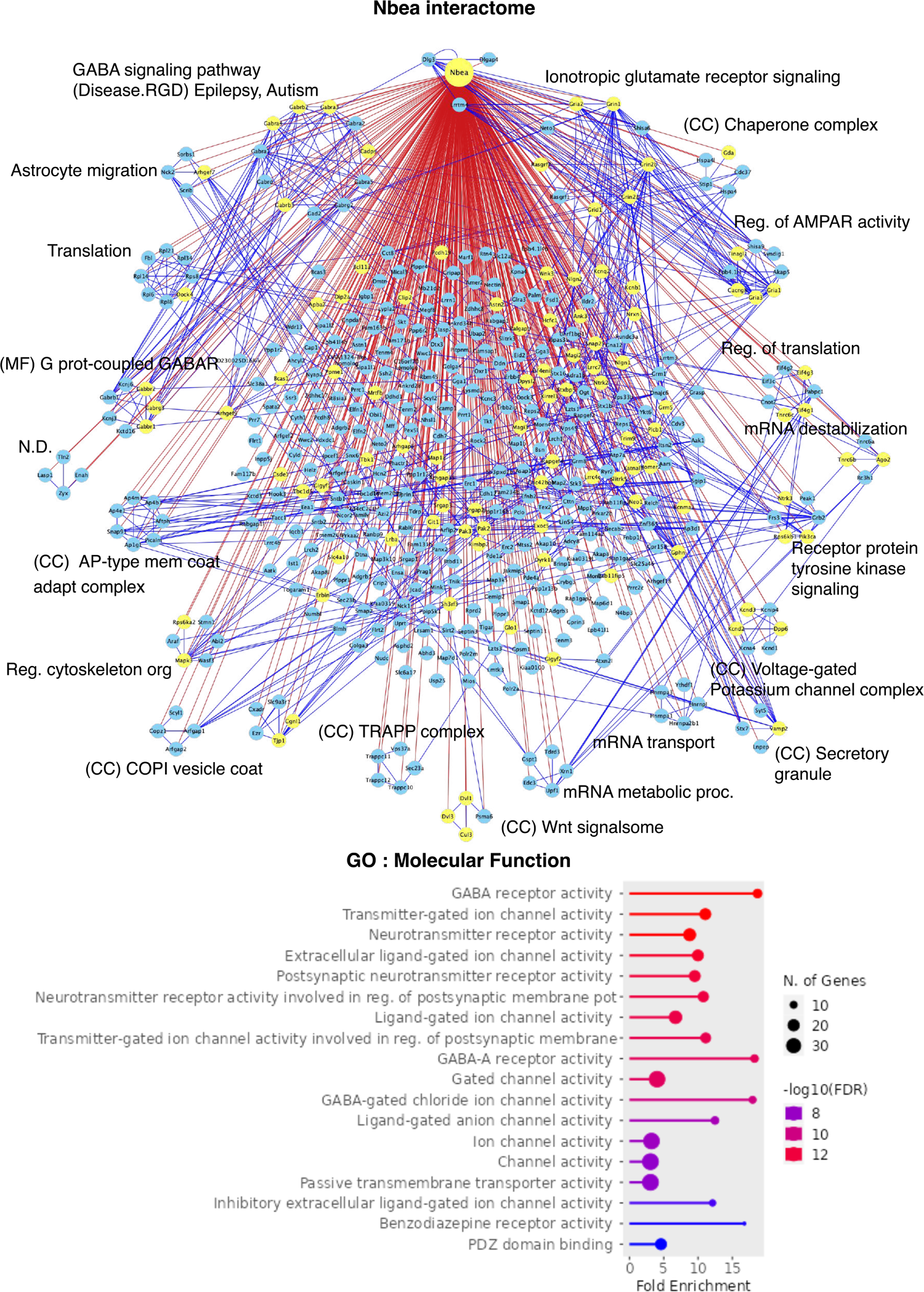
Individual interactome networks of 14 autism risk proteins and the additional 2 “hub” proteins. Interactome network associated with the bait protein is shown in each Figure. Blue lines denote STRING interactions and red lines signify identified HiUGE-iBioID interactions. Yellow nodes highlight proteins encoded by SFARI gene orthologs. Annotations denote exemplary significant gene ontology (GO) terms associated with the protein clusters segregated by MCL. Unless otherwise specified, the Biological Process pathway database was used. CC: Cellular Component pathway database. Charts of GO results of the overall bait interactomes using the Molecular Function (MF) pathway database are shown as well, below each network plot.

**Fig. S17.**
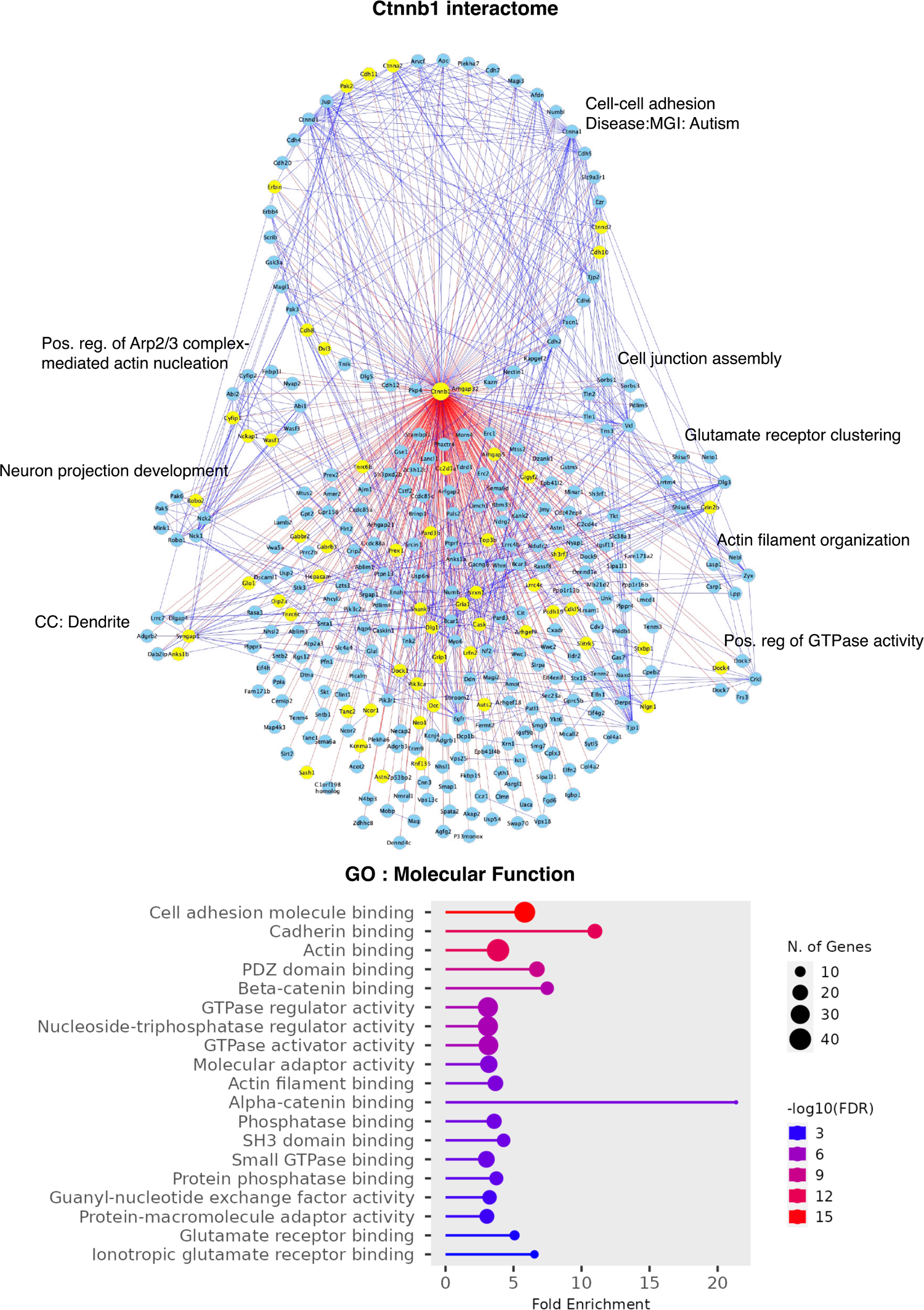
Individual interactome networks of 14 autism risk proteins and the additional 2 “hub” proteins. Interactome network associated with the bait protein is shown in each Figure. Blue lines denote STRING interactions and red lines signify identified HiUGE-iBioID interactions. Yellow nodes highlight proteins encoded by SFARI gene orthologs. Annotations denote exemplary significant gene ontology (GO) terms associated with the protein clusters segregated by MCL. Unless otherwise specified, the Biological Process pathway database was used. CC: Cellular Component pathway database. Charts of GO results of the overall bait interactomes using the Molecular Function (MF) pathway database are shown as well, below each network plot.

**Fig. S18.**
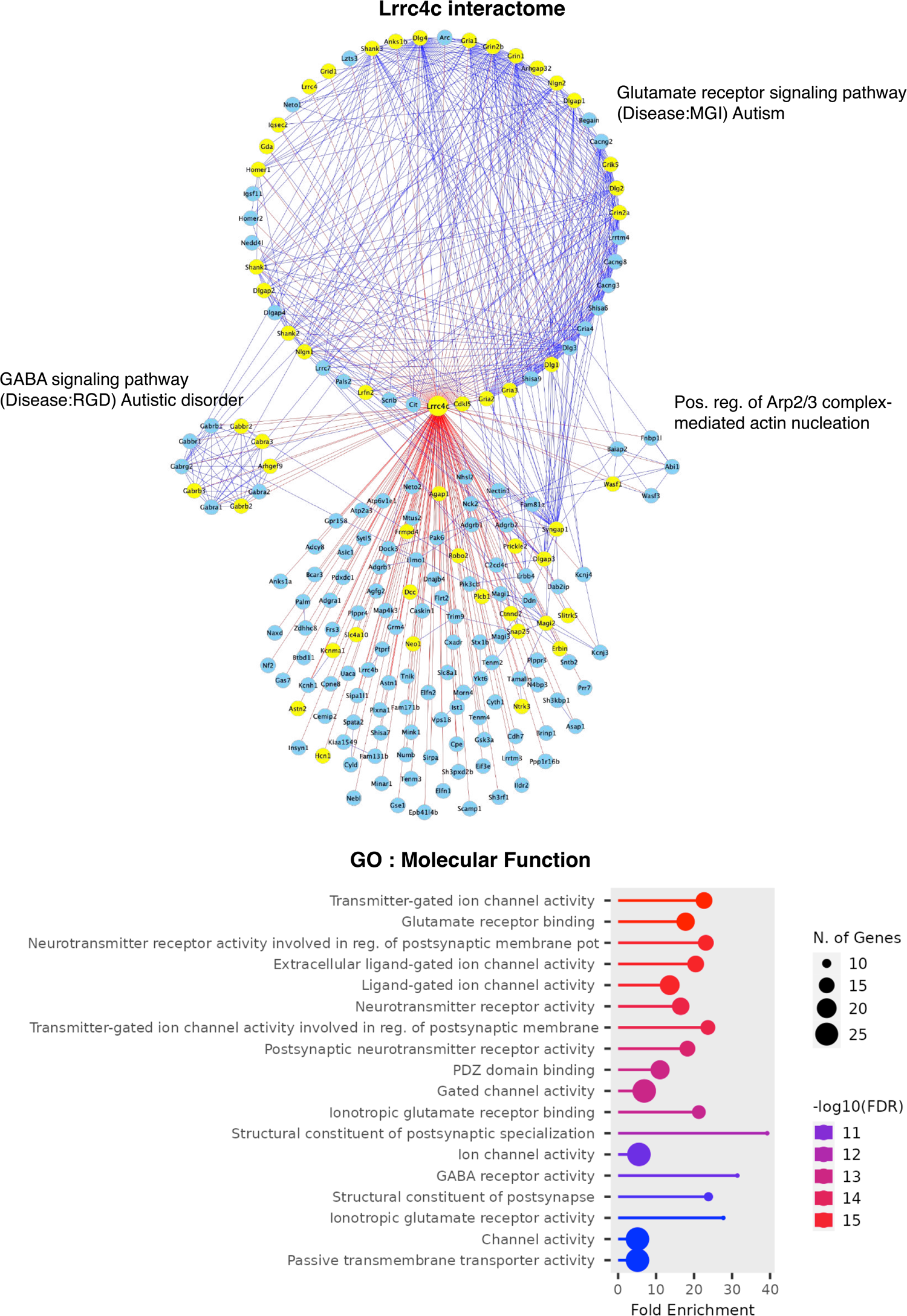
Individual interactome networks of 14 autism risk proteins and the additional 2 “hub” proteins. Interactome network associated with the bait protein is shown in each Figure. Blue lines denote STRING interactions and red lines signify identified HiUGE-iBioID interactions. Yellow nodes highlight proteins encoded by SFARI gene orthologs. Annotations denote exemplary significant gene ontology (GO) terms associated with the protein clusters segregated by MCL. Unless otherwise specified, the Biological Process pathway database was used. CC: Cellular Component pathway database. Charts of GO results of the overall bait interactomes using the Molecular Function (MF) pathway database are shown as well, below each network plot.

**Fig. S19.**
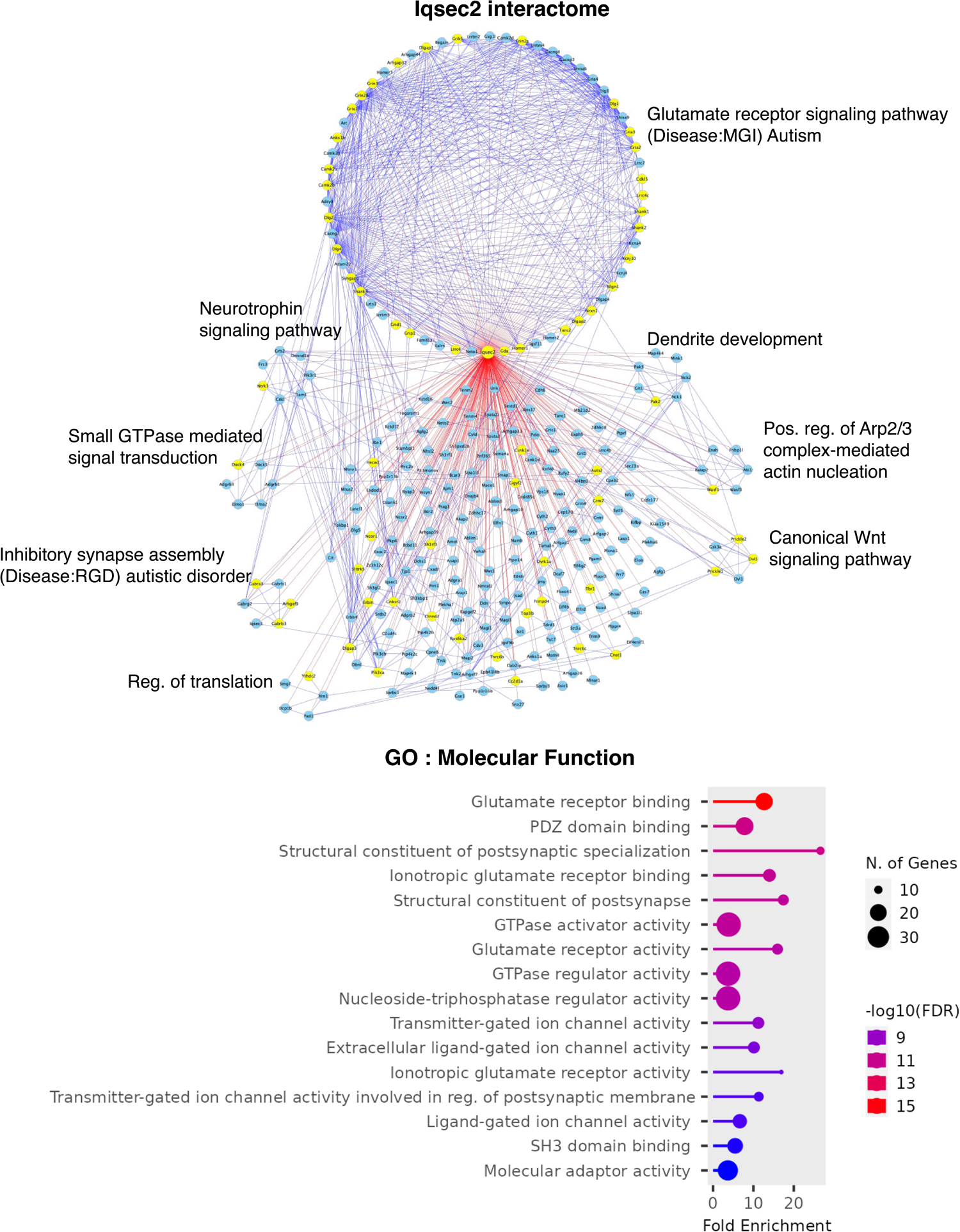
Individual interactome networks of 14 autism risk proteins and the additional 2 “hub” proteins. Interactome network associated with the bait protein is shown in each Figure. Blue lines denote STRING interactions and red lines signify identified HiUGE-iBioID interactions. Yellow nodes highlight proteins encoded by SFARI gene orthologs. Annotations denote exemplary significant gene ontology (GO) terms associated with the protein clusters segregated by MCL. Unless otherwise specified, the Biological Process pathway database was used. CC: Cellular Component pathway database. Charts of GO results of the overall bait interactomes using the Molecular Function (MF) pathway database are shown as well, below each network plot.

**Fig. S20.**
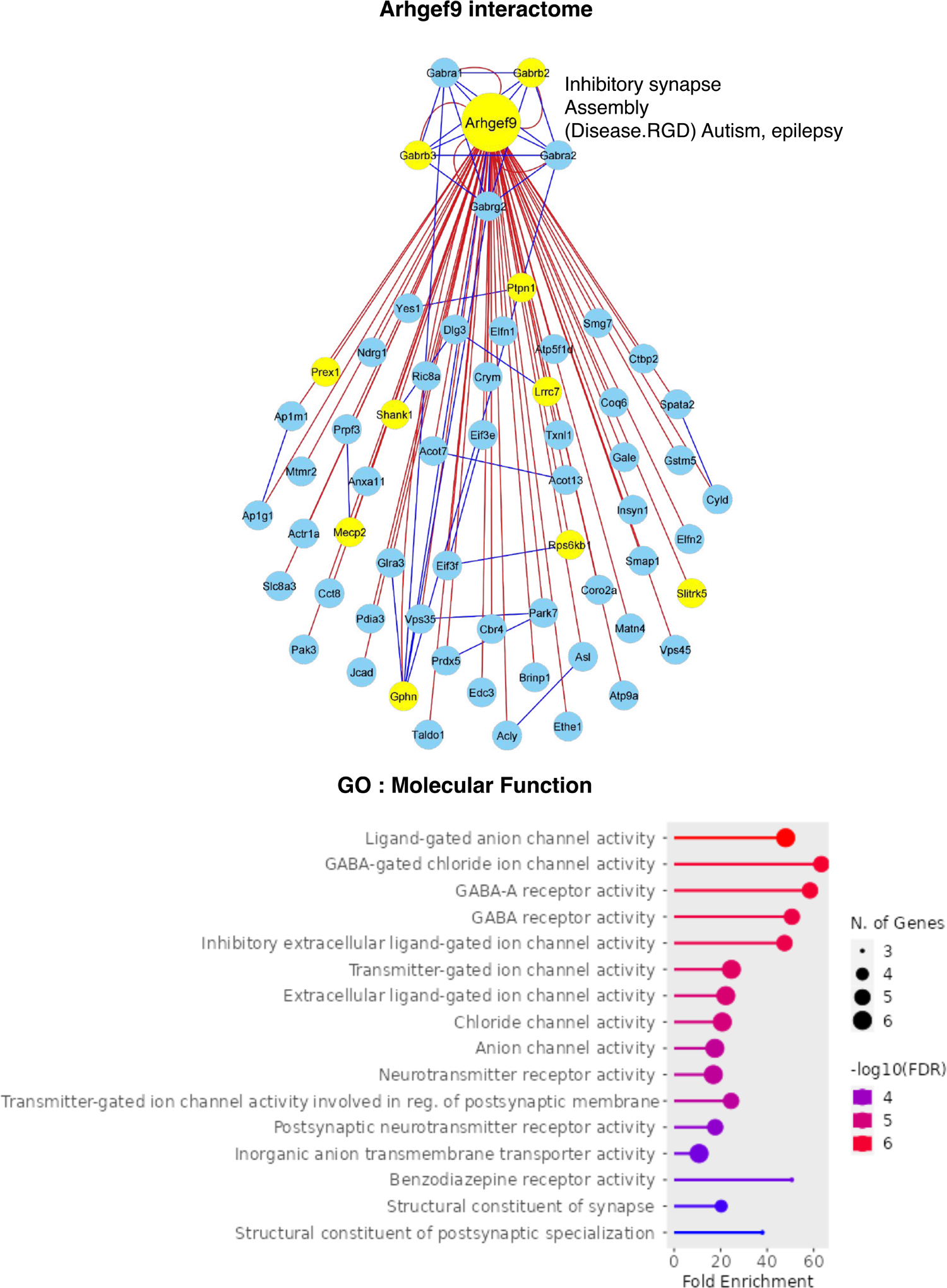
Individual interactome networks of 14 autism risk proteins and the additional 2 “hub” proteins. Interactome network associated with the bait protein is shown in each Figure. Blue lines denote STRING interactions and red lines signify identified HiUGE-iBioID interactions. Yellow nodes highlight proteins encoded by SFARI gene orthologs. Annotations denote exemplary significant gene ontology (GO) terms associated with the protein clusters segregated by MCL. Unless otherwise specified, the Biological Process pathway database was used. CC: Cellular Component pathway database. Charts of GO results of the overall bait interactomes using the Molecular Function (MF) pathway database are shown as well, below each network plot.

**Fig. S21.**
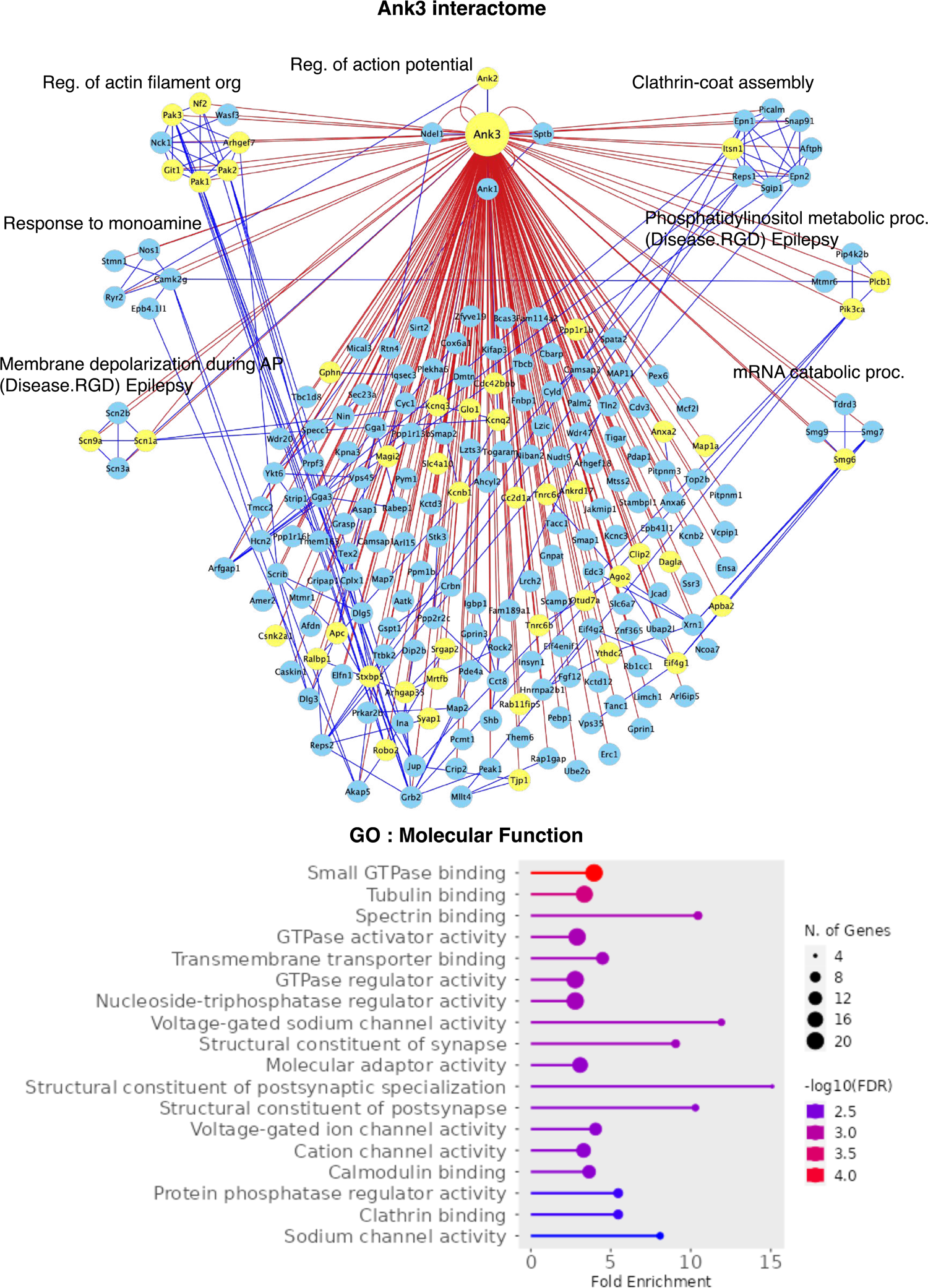
Individual interactome networks of 14 autism risk proteins and the additional 2 “hub” proteins. Interactome network associated with the bait protein is shown in each Figure. Blue lines denote STRING interactions and red lines signify identified HiUGE-iBioID interactions. Yellow nodes highlight proteins encoded by SFARI gene orthologs. Annotations denote exemplary significant gene ontology (GO) terms associated with the protein clusters segregated by MCL. Unless otherwise specified, the Biological Process pathway database was used. CC: Cellular Component pathway database. Charts of GO results of the overall bait interactomes using the Molecular Function (MF) pathway database are shown as well, below each network plot.

**Fig. S22.**
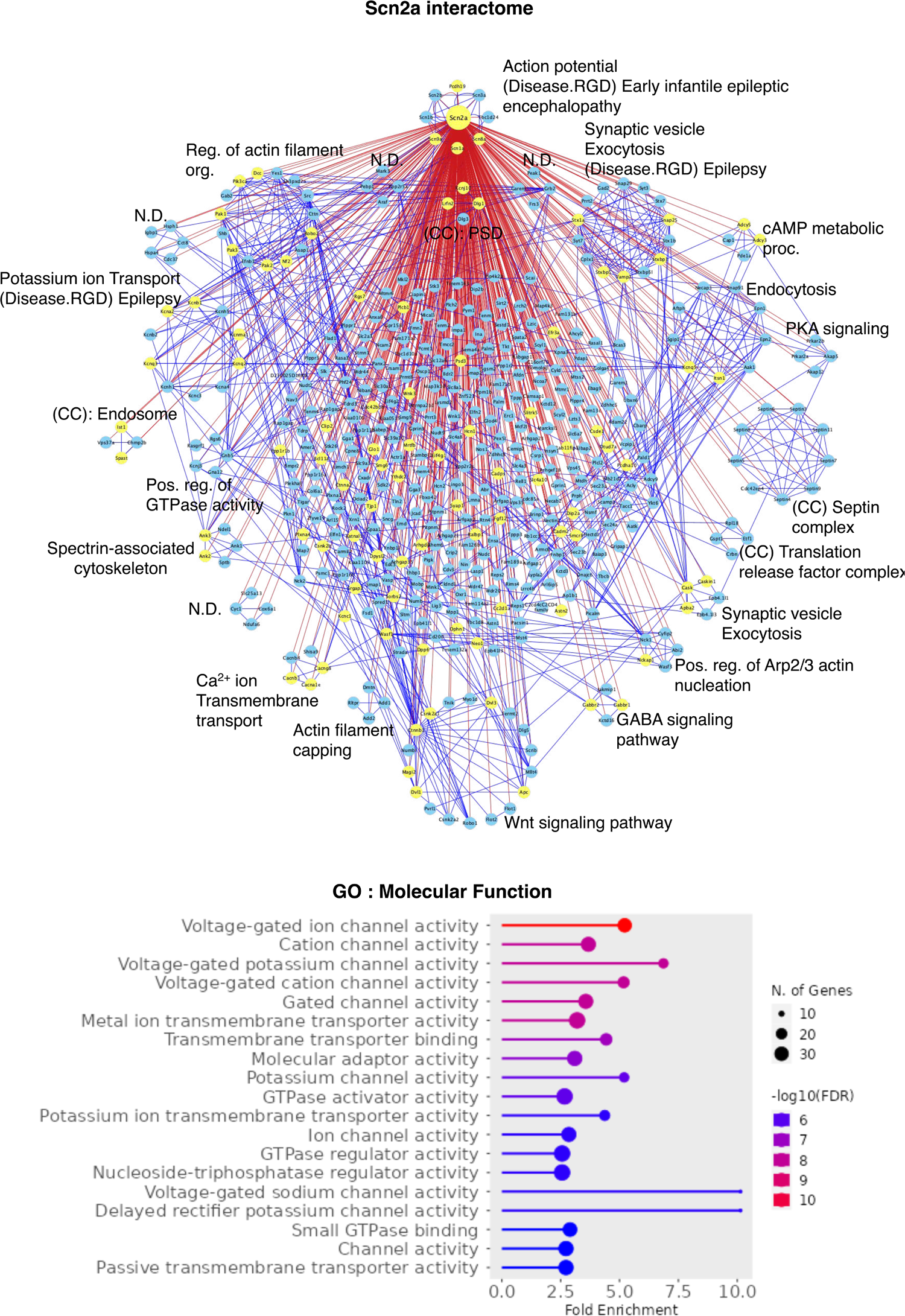
Individual interactome networks of 14 autism risk proteins and the additional 2 “hub” proteins. Interactome network associated with the bait protein is shown in each Figure. Blue lines denote STRING interactions and red lines signify identified HiUGE-iBioID interactions. Yellow nodes highlight proteins encoded by SFARI gene orthologs. Annotations denote exemplary significant gene ontology (GO) terms associated with the protein clusters segregated by MCL. Unless otherwise specified, the Biological Process pathway database was used. CC: Cellular Component pathway database. Charts of GO results of the overall bait interactomes using the Molecular Function (MF) pathway database are shown as well, below each network plot.

**Fig. S23.**
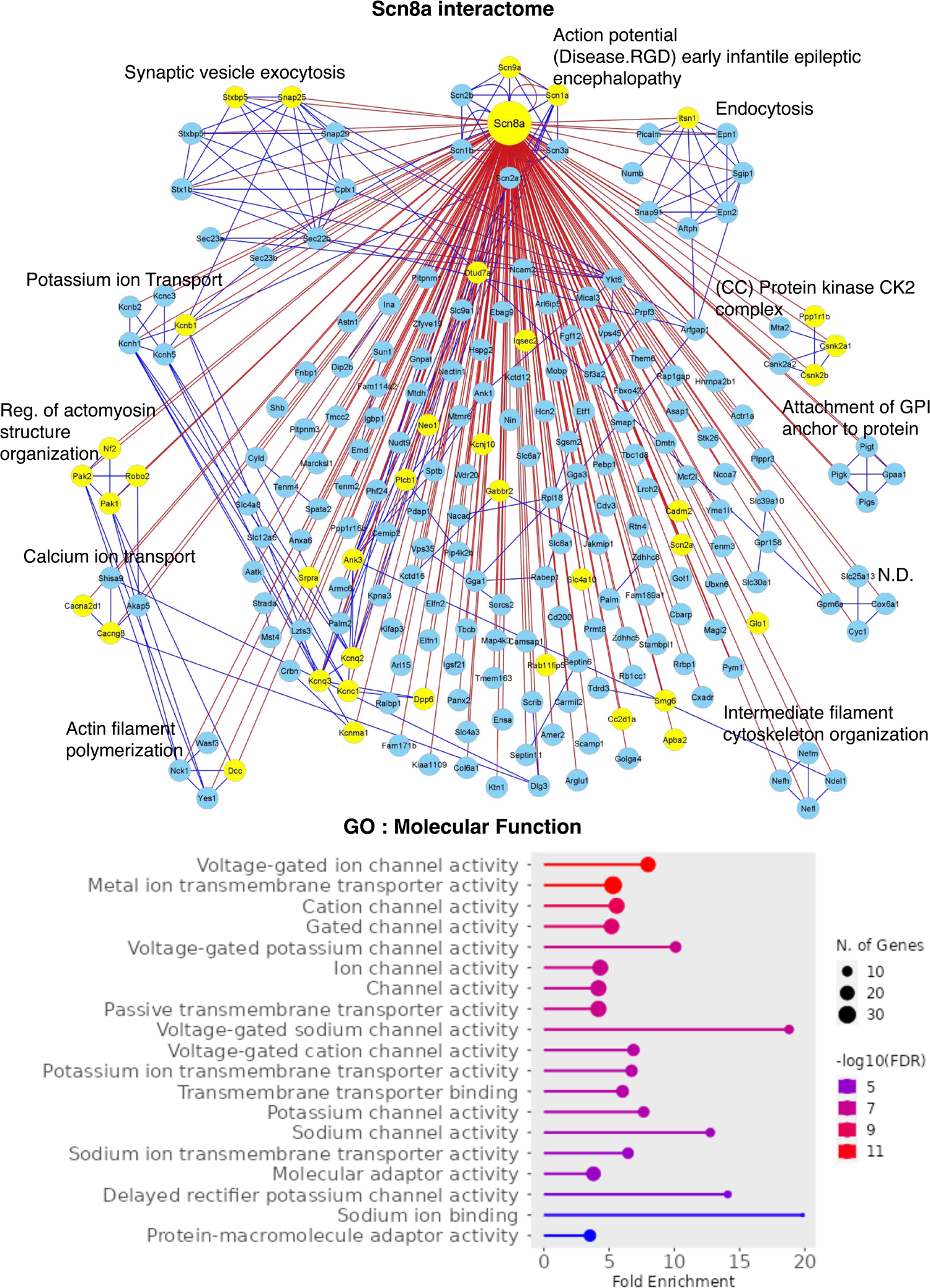
Individual interactome networks of 14 autism risk proteins and the additional 2 “hub” proteins. Interactome network associated with the bait protein is shown in each Figure. Blue lines denote STRING interactions and red lines signify identified HiUGE-iBioID interactions. Yellow nodes highlight proteins encoded by SFARI gene orthologs. Annotations denote exemplary significant gene ontology (GO) terms associated with the protein clusters segregated by MCL. Unless otherwise specified, the Biological Process pathway database was used. CC: Cellular Component pathway database. Charts of GO results of the overall bait interactomes using the Molecular Function (MF) pathway database are shown as well, below each network plot.

**Fig. S24.**
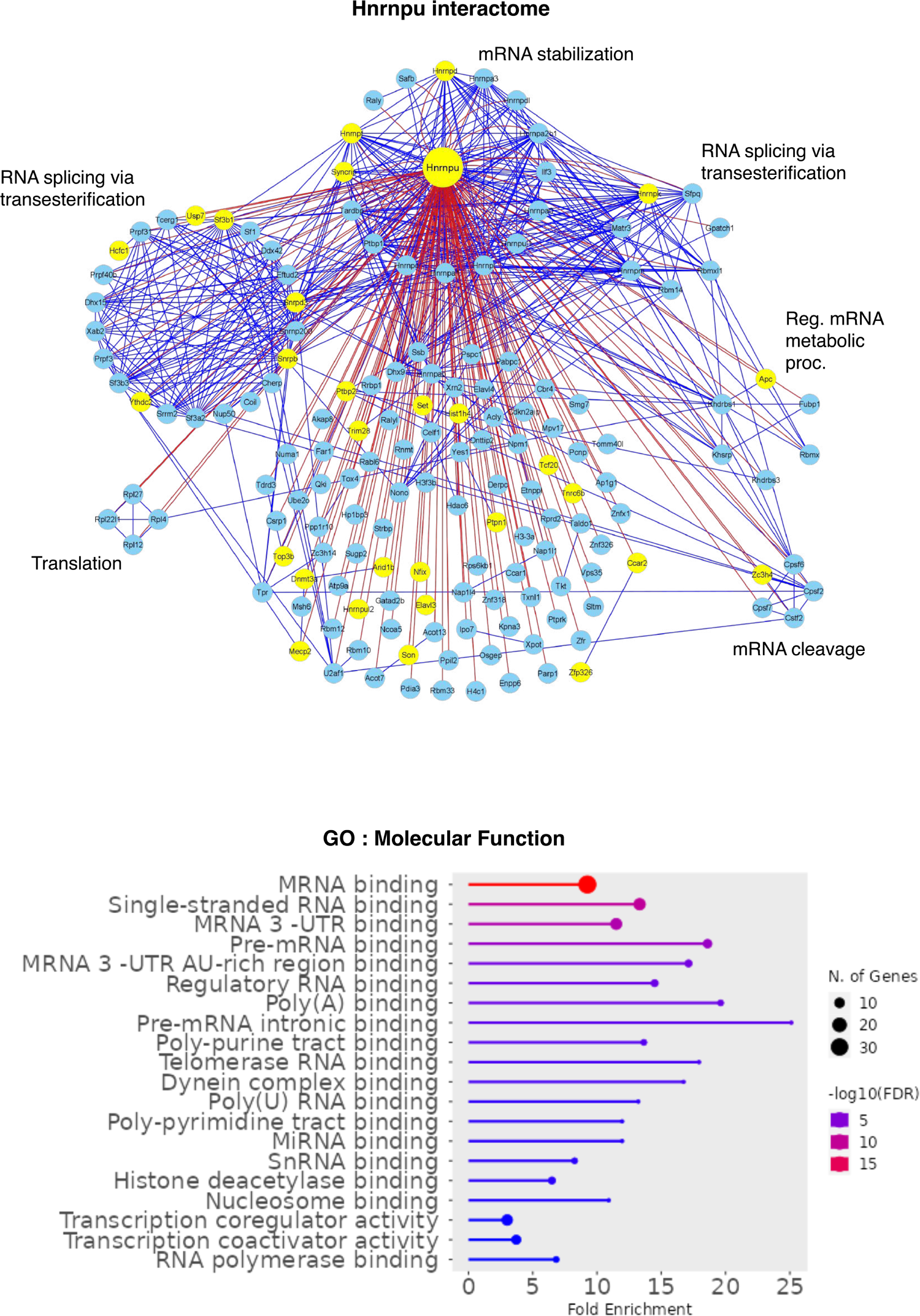
Individual interactome networks of 14 autism risk proteins and the additional 2 “hub” proteins. Interactome network associated with the bait protein is shown in each Figure. Blue lines denote STRING interactions and red lines signify identified HiUGE-iBioID interactions. Yellow nodes highlight proteins encoded by SFARI gene orthologs. Annotations denote exemplary significant gene ontology (GO) terms associated with the protein clusters segregated by MCL. Unless otherwise specified, the Biological Process pathway database was used. CC: Cellular Component pathway database. Charts of GO results of the overall bait interactomes using the Molecular Function (MF) pathway database are shown as well, below each network plot.

**Fig. S25.**
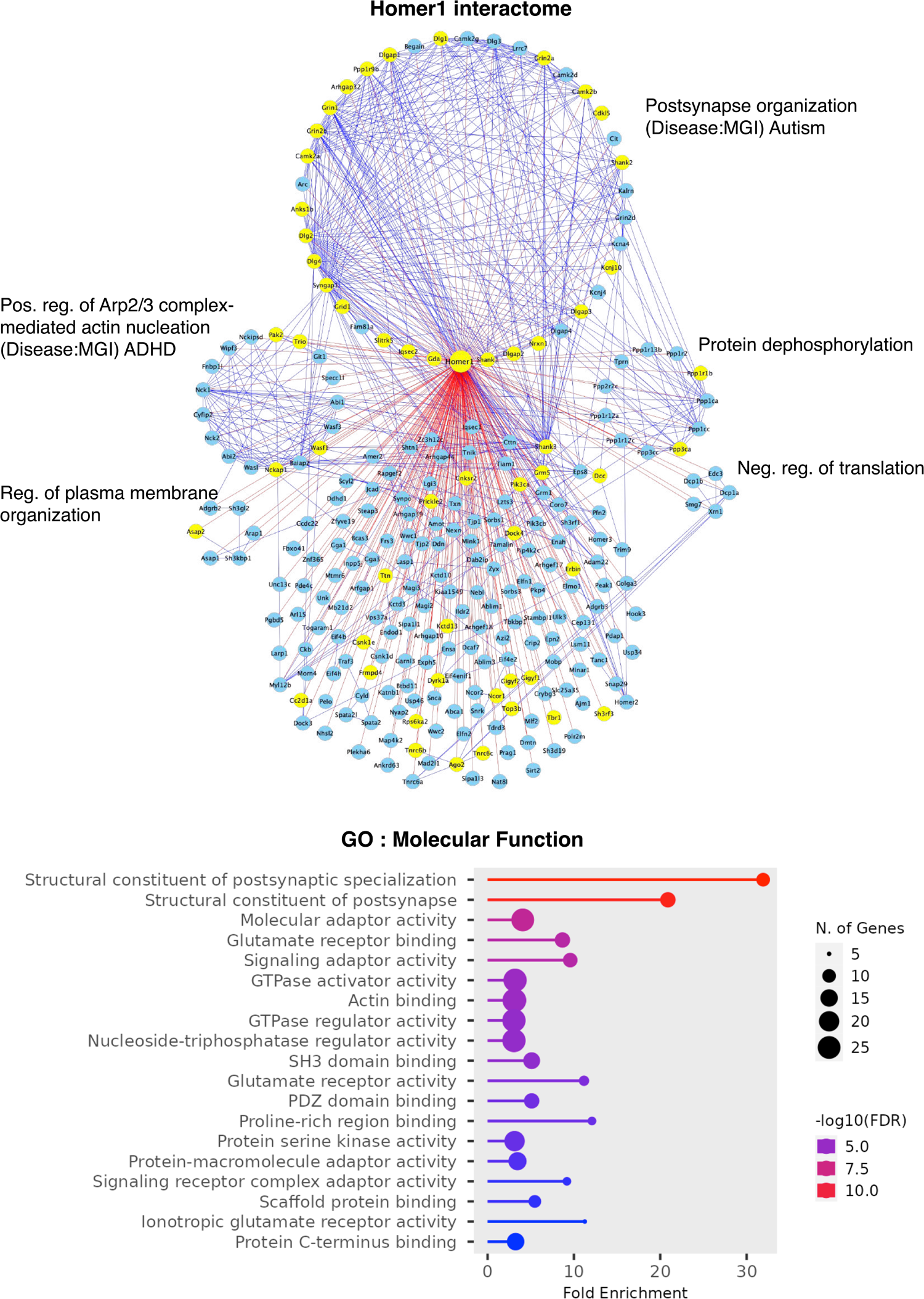
Individual interactome networks of 14 autism risk proteins and the additional 2 “hub” proteins. Interactome network associated with the bait protein is shown in each Figure. Blue lines denote STRING interactions and red lines signify identified HiUGE-iBioID interactions. Yellow nodes highlight proteins encoded by SFARI gene orthologs. Annotations denote exemplary significant gene ontology (GO) terms associated with the protein clusters segregated by MCL. Unless otherwise specified, the Biological Process pathway database was used. CC: Cellular Component pathway database. Charts of GO results of the overall bait interactomes using the Molecular Function (MF) pathway database are shown as well, below each network plot.

**Fig. S26.**
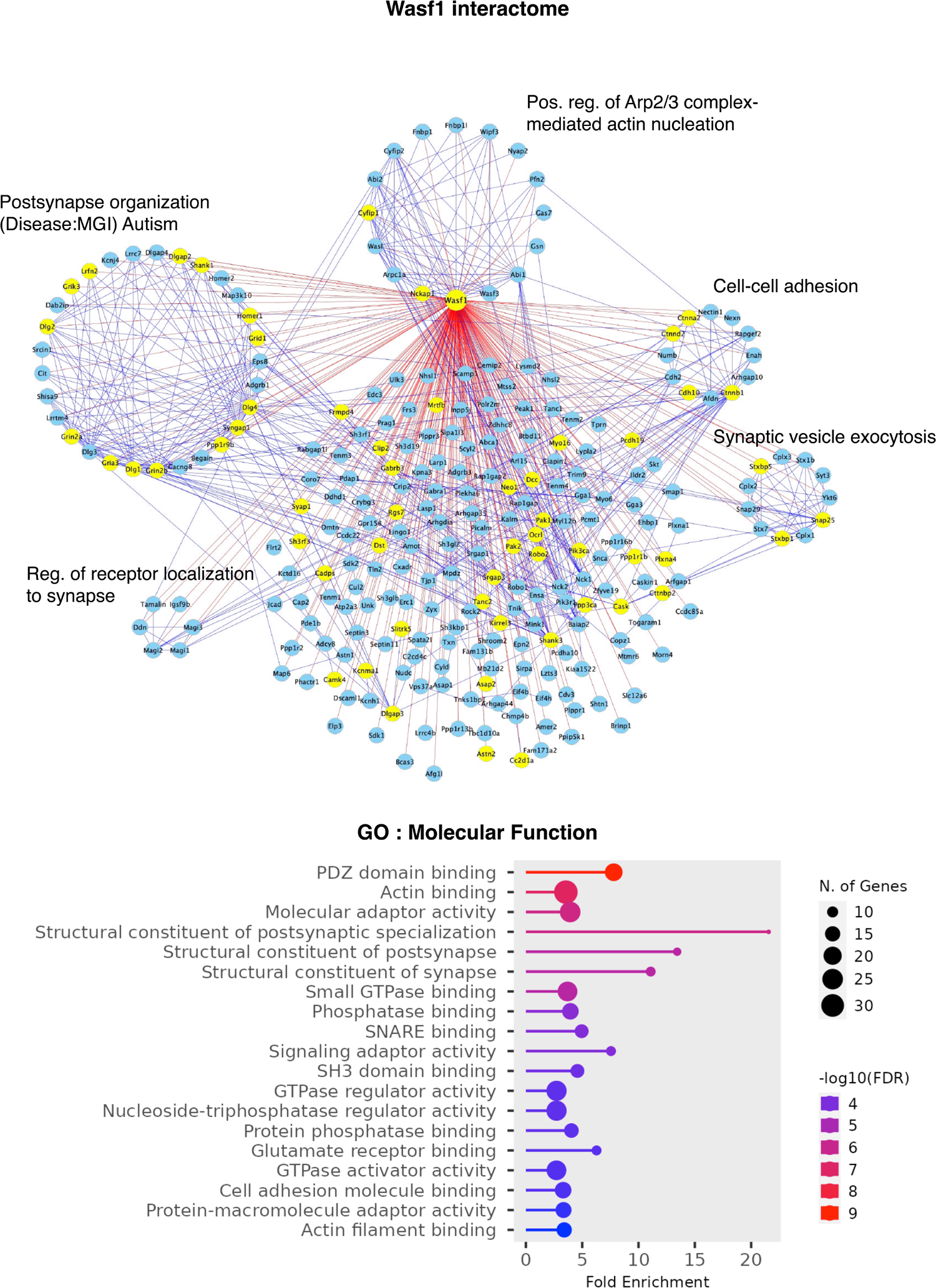
Individual interactome networks of 14 autism risk proteins and the additional 2 “hub” proteins. Interactome network associated with the bait protein is shown in each Figure. Blue lines denote STRING interactions and red lines signify identified HiUGE-iBioID interactions. Yellow nodes highlight proteins encoded by SFARI gene orthologs. Annotations denote exemplary significant gene ontology (GO) terms associated with the protein clusters segregated by MCL. Unless otherwise specified, the Biological Process pathway database was used. CC: Cellular Component pathway database. Charts of GO results of the overall bait interactomes using the Molecular Function (MF) pathway database are shown as well, below each network plot.

**Fig. S27.**
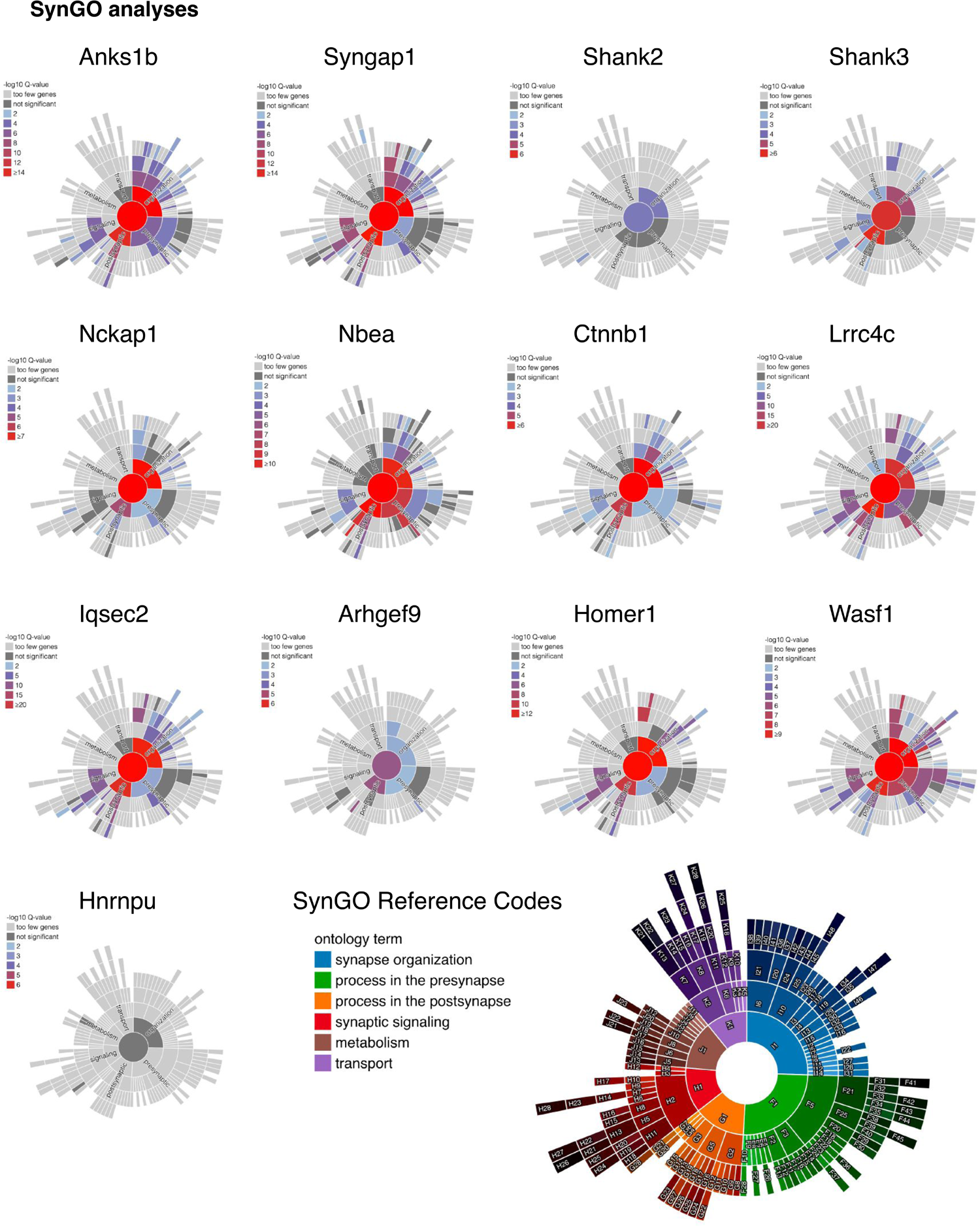
SynGO analyses of the synaptic bait interactomes versus the nucleus bait interactome. SynGO analyses showing the expected functions of the synaptic bait interactomes mainly within the domains of “synapse organization” and “process in the postsynapse”. In contrast, no significant SynGO enrichment was detected for the nucleus Hnrnpu interactome.

**Table S1. Proteomic results and statistics of HiUGE-iBioID experiments.**

Proteomic results and statistics of HiUGE-iBioID experiments are shown. Batch-1: Anks1b, Shank2, Nckap1; Batch-2: Nbea; Batch-3: Ank3, Scn2a, Scn8a; Batch-4: Arhgef9, Hnrnpu; Batch-5: Shank3; Batch-6: Syngap1-Trunc; Batch-7: Soluble TurboID; Batch-8: Syngap1, Ctnnb1, Iqsec2, Lrrc4c (modified strategies to preserve PDZ-binding motifs); Batch-9: Homer1, Wasf1.

**Table S2. HiUGE-iBioID interactomes and their overlaps with autism gene lists.**

Filtered gene lists of the detected interactomes associated with 14 high-risk autism targets are shown. The genes that overlap with the SFARI database, Satterstrom et al. ^3^, and Fu et al. ^7^ gene lists are shown in a separate tab.

**Table S3. Proteomic detection following antibody immunoprecipitations.**

Proteomic results and statistics of Anks1b and Scn2a immunoprecipitation experiments.

**Table S4. Proteomic results and statistics of spatial co-perturbation experiments.**

Proteomic results and statistics of spatial co-perturbation experiments are shown, including comparisons of Syngap1-Het synaptosome proteome with WT in the cortex and striatum, and comparison of Scn2a^+/R102Q^ LOPIT-DC fraction-5 proteome with WT.

**Table S5. Statistical domain for GO analyses.**

Statistical domain for GO analyses compiled from cumulative proteomic detections of brain-derived samples in our lab.

**Table S6. Statistics for behavioral characterization of Scn2a^+/R102Q^ mice.**

Metrics and statistic summaries for the elevated zero maze, hole-board, self-grooming, USV, resident-intruder, and social dyadic tests are reported in this Table. For the USV test, no statistically significant differences were detected in the following pre-social (baseline) responses: Number of calls: WT (8.0 ± 1.6), Scn2a^+/R102Q^ (5.9 ± 2.8) [t(22)=0.693, *p*=0.496]; and Call frequency (kHz): WT (30.4 ± 2.7), Scn2a^+/R102Q^ (20.0 ± 5.5) [t(22)=1.858, *p*=0.077]. Difference in call duration (msec) was marginal: WT (6.8 ± 0.8), Scn2a^+/R102Q^ (4.0 ± 1.2) [t(22)=2.070, *p*=0.050]; n = 14 WT males, n = 10 Scn2a^+/R102Q^ males; data presented as means ± SEM, independent samples t-tests, two-tailed.

**Table S7. Results of the enrichment analyses between the HiUGE-iBioID interactomes and an autism GWAS study.**

Results of the enrichment analyses between the HiUGE-iBioID and Grove et al., 2019, autism genome-wide association meta-analysis (GWAS) using MAGMA.

